# Temporal TOC1-CLV3-WUS cascade controls organogenesis and rhythmic stem development

**DOI:** 10.64898/2026.09.24.753998

**Authors:** Jiahong Qi, Jiahui Xu, Yufeng Yang, Yichao Zheng, Mei Jin, Lin Guo, Xiaodong Xu, Qiguang Xie, Hao Zhang, Xigang Liu

**Author notes:** **Corresponding authors Email:** (H.Z.), (X.L.). These authors contributed equally to this work (Jiahong Qi, Jiahui Xu, Yufeng Yang).

## Abstract

Coordinated organogenesis and rhythmic stem development determine shoot morphogenesis, which governs distinct plant architecture and crop yield. The shoot apical meristem (SAM) drives organ initiation and stem growth, with its activity sustained by the spatially defined *CLAVATA3* (*CLV3*)-*WUSCHEL* (*WUS*) feedback loop. However, how the temporal *CLV3*-*WUS* module modulates SAM activity remains elusive. Here, we demonstrate that both *CLV3* and *WUS* are required for circadian stem elongation. The clock component TIMING OF CAB EXPRESSION1 (TOC1) directly activates and sustains the circadian expression of *CLV3*, and this regulatory process is controlled by the *APETALA1*-*CIRCADIAN CLOCK-ASSOCIATED1* module. Notably, the rhythmic expression pattern of *CLV3*, rather than its transcript abundance, plays a more important role in regulating *WUS* expression and sustaining SAM activity, and CLV3 confers circadian rhythmicity to WUS protein accumulation. Transcriptome and ChIP seq analyses reveal that WUS modulates multiple developmental programs, including meristem homeostasis, phytohormone responses, and cell wall remodeling. Mechanistically, WUS directly activates *PECTIN METHYLESTERASE5* expression to promote stem elongation. Collectively, our findings uncover a spatiotemporal regulatory cascade that fine-tunes SAM activity to coordinate rhythmic organogenesis and stem development, thereby establishing optimal phyllotactic patterning in plants.

## INTRODUCTION

The three-dimensional architecture of flowering plants is established through the continuous and coordinated production of lateral organs and axial stem tissues from the shoot apical meristem (SAM). As the ultimate source of all above-ground organs, the SAM maintains a pool of pluripotent stem cells for the initiation of leaves, flowers and the intervening stem internodes^1–3^. This dual output, organogenesis and stem elongation, must be precisely coordinated in space and time to generate optimal shoot architectures that determine light interception, photosynthetic efficiency and, ultimately, crop yield ^1,4,5^. Indeed, the modification of plant architecture, particularly stem height and phyllotaxis, was central to the Green Revolution, and continues to offer untapped potential for the next generation of sustainable crop improvement ^6–8^. Despite its fundamental importance, however, the molecular mechanisms that coordinate organogenesis with stem development have received far less attention than the study of individual organs such as leaves or flowers ^2^.

The SAM is spatially organized into functionally distinct domains: the central zone (CZ) harbours the stem cell population; the peripheral zone (PZ) gives rise to organ primordia; and the rib zone (RZ), located beneath the CZ, contains the rib meristem (RM) that provides the cellular basis for stem axial growth ^2,3,9,10^. Stem cell homeostasis in the SAM is governed by a well-characterized negative feedback loop between the homeodomain transcription factor WUSCHEL (WUS) and the CLAVATA (CLV) class of gene *CLV3* ^11–13^. *WUS*, expressed in the organizing centre (OC) beneath the stem cells, moves non-cell-autonomously to the CZ, where it activates *CLV3* transcription ^14,15^. The mature CLV3 peptide is secreted into the apoplast to restrict *WUS* expression both transcriptionally and post-translationally and thereby maintaining meristem size and organogenic capacity ^12–14,16,17^. This spatial regulatory circuit has been established as a central paradigm for plant stem cell maintenance ^11–13,18^. While this spatial framework has been extensively characterized, whether and how the *CLV3*-*WUS* circuit operates under temporal control to coordinate organogenesis with stem elongation has remained largely unexplored.

Rhythmic stem elongation is particularly evident in Arabidopsis inflorescence stems, where growth occurs predominantly at night and is tightly coupled with the periodic initiation of organ primordia to establish optimal shoot architecture^1–3,19^. Notably, the characteristic Fibonacci spiral patterns observed in most land plants depend on the precise temporal spacing of organ initiation relative to stem growth ^20–22^, highlighting the intimate relationship between time and form in plant development. The *CLV3*-*WUS* module exhibits striking developmental phenotypes when disrupted: *clv3* mutants produce enlarged meristems with supernumerary organs, whereas *wus* mutants undergo premature meristem termination^4,11–13^, raising the possibility that this module contributes not only to spatial meristem organization but also to the temporal gating of growth. However, whether and how the circadian clock interfaces with the *CLV3*-*WUS* circuitry to temporally coordinate organogenesis and stem elongation has remained an open question.

The circadian clock is an endogenous timekeeping mechanism that synchronizes diverse physiological and developmental processes with the diurnal environment ^23^. By integrating cyclic environmental signals such as light and temperature, the clock embeds temporal information into the plant’s developmental program, conferring adaptive fitness advantages ^24–26^. In plants, the SAM has been proposed to function analogously to the suprachiasmatic nucleus in mammals, acting as a central pacemaker that coordinates rhythmic outputs to distal organs such as roots and leaves ^27,28^. Core clock components form interlocked transcriptional–translational feedback loops that generate ∼24-h oscillations in gene expression ^29,30^. Among these, *TIMING OF CAB EXPRESSION1* (*TOC1*) encodes an evening-expressed pseudo-response regulator that has been characterized predominantly as a transcriptional repressor, binding to Evening Element (EE) motifs in target promoters to repress morning-phase genes such as *CIRCADIAN CLOCK ASSOCIATED1*(*CCA1*) and *LATE ELONGATED HYPOCOTYL* (*LHY*) ^31–33^. Recent evidence, however, has revealed that TOC1 can also function as a transcriptional activator in specific spatiotemporal contexts, through interaction with NF-Y complexes and recruitment of tissue-specific co-factors ^34^. In rice, OsTOC1 has been shown to act as both a repressor and an activator of circadian outputs, suggesting that the regulatory repertoire of TOC1 is broader than previously appreciated ^35^. We recently found that TOC1 acts downstream of AP1 to non-cell-autonomously regulate rhythmic stem elongation by promoting expression of *VANGUARD1* (*VGD1*), one of *Pectin Methylesterases* (*PME*) genes ^36^, yet the downstream effector through which TOC1 temporally coordinates organogenesis with stem growth has remained unclear.

Stem elongation ultimately depends on the extensibility of the cell wall, a dynamic structure whose mechanical properties are largely determined by the degree of pectin methylesterification ^37^. PMEs catalyse the demethylesterification of homogalacturonan, exposing negatively charged carboxyl groups that modulate calcium cross-linking and cell wall stiffness, thereby controlling cell elongation and division ^38–40^. During inflorescence stem growth, spatiotemporal gradients of PME activity correlate closely with cell wall mechanical gradients, and specific *PME* genes, such as *PECTIN METHYLESTERASE5* (*PME5*) and *VGD1*, have been implicated in cell expansion ^40–43^. Remarkably, recent studies have demonstrated that WUS directly regulates the expression of a subset of *PME* genes in the SAM, thereby sustaining the mechanical properties of stem cell walls and coordinating organogenesis ^43,44^. These findings place PME-mediated cell wall remodeling as a downstream executive mechanism of the *CLV3*-*WUS* regulatory network.

Here, we uncover a spatiotemporal regulatory cascade in which the circadian clock, through the AP1/CCA1-TOC1 module, gates the rhythmic expression of *CLV3* to coordinate organogenesis and circadian stem elongation. We demonstrate that TOC1 directly activates *CLV3* transcription, sustaining its evening-phased circadian expression, and that this temporal pattern, rather than the absolute transcript level, is critical for maintaining *CLV3*-*WUS* feedback loop function and SAM homeostasis. We further show that CLV3 confers circadian rhythmicity to WUS protein accumulation, and that WUS directly activates *PME5* expression to promote stem elongation.

## RESULTS

### *CLV3* and *WUS* are required for the normal organogenesis and rhythmic stem elongation

To assess the roles of *CLV3* and *WUS* in stem development, we examined the stem growth of wild type (WT), *clv3-1* (one of null mutants of *CLV3*) ^45^ and *wus-7* (one weak allele of *wus* mutants ^46^, since null mutant of *wus* only generates bushy shoots) under normal condition (16h light/8h dark). At 28 days after germination (DAG), *clv3-1* exhibited moderately longer stems compared with the WT, suggesting that loss-of-function of *CLV3* promotes stem elongation. By contrast, *wus-7* displayed drastically shortened stems relative to WT. Together with the bushy-shoot phenotype reported for *wus-1* (a null mutant of *WUS*) ^47^, these observations demonstrate that *WUS* is indispensable for stem initiation and elongation (Fig. 1a and Extended Data Fig. 1a). Since differences in flowering time could affect stem elongation and organogenesis, we further examined the transition from vegetative to reproductive growth in these three genotypes. The number of rosette leaves, a general indicator of flowering time at bolting, was significantly increased in *clv3-1*, whereas it was slightly reduced in *wus-7* compared with the WT (Extended Data Fig. 1b). However, the rosette leaf number is also influenced by meristem activity, we then performed histological sectioning of shoot apices at 9, 11, 13, and 15 DAG to more precisely determine the floral-transition timing. Floral primordia were first observed at 11 DAG and became prominent at 13 DAG in *clv3-1*. By contrast, floral primordia emerged at 13 DAG in both L*er* and *wus-7* plants (Extended Data Fig. 1c). Notably, although *wus-7* underwent floral transition at a time comparable to the WT, it exhibited severe defects in stem growth and silique production. These results suggested that both *CLV3* and *WUS* were required for the stem elongation.

**Fig. 1.**
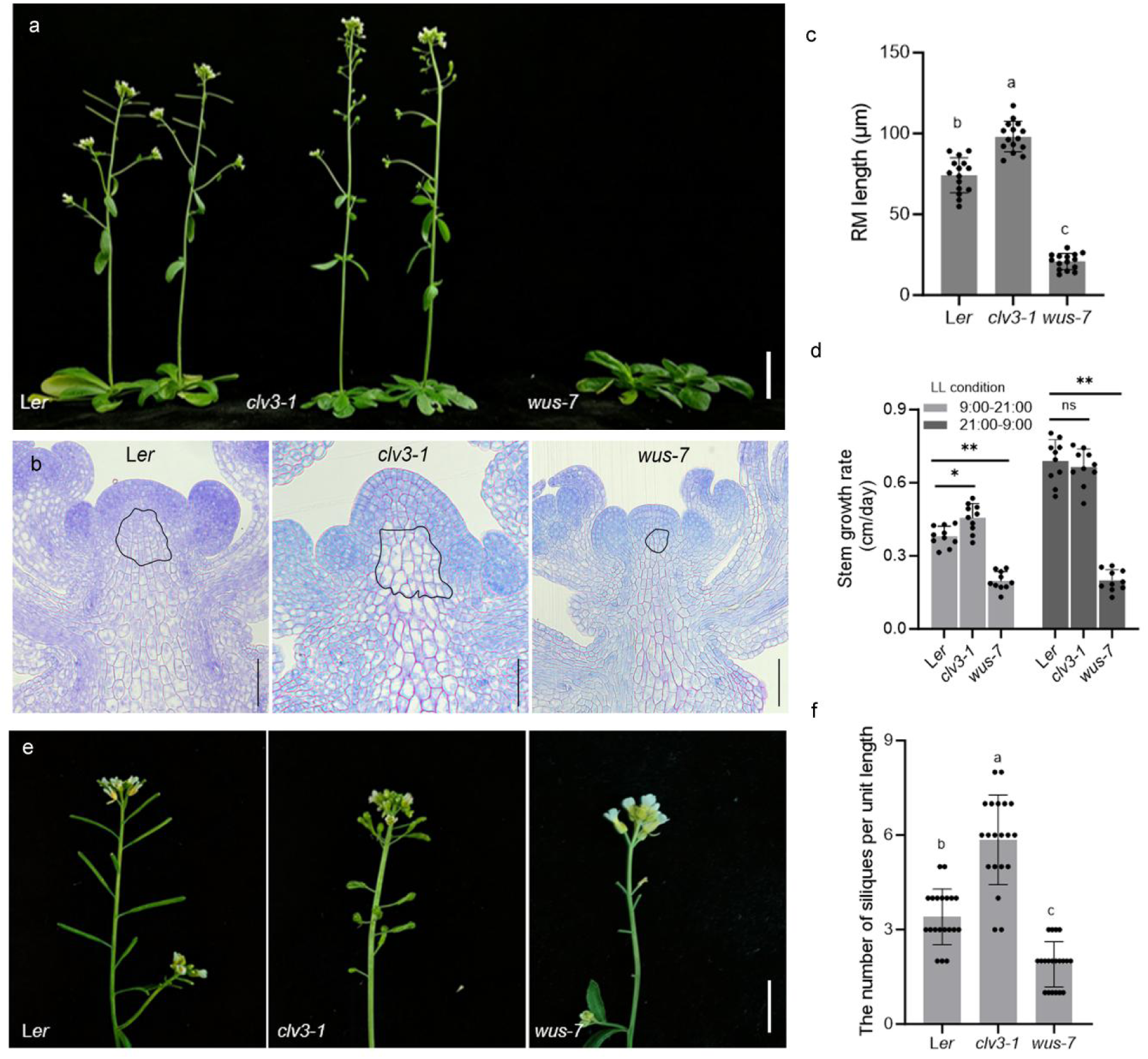
CLV3 and WUS are required for normal organogenesis and rhythmic stem elongation. (**a**) Morphological observation of L*er*, *clv3-1* and *wus-7* plants. Scale bar: 2 cm. (**b**) Longitudinal sections of shoot apices of L*er*, *clv3-1* and *wus-7* showing the Rib Meristem (RM) region outlined by black lines. Scale bars = 50 μm. (**c**) Statistical analysis of the length of RM of indicated plants. Data represent mean ± SD (*n =* 15). Different letters indicate statistically significant differences by one-way ANOVA (Tukey’s multiple comparisons test, *P <* 0.01). (**d**) The stem growth rate of indicated plants under LL (constant-light) conditions during subjective day and night. 9:00–21:00, subjective day; 21:00–9:00, subjective night. Data represent mean ± SD (*n* = 10). Asterisks indicate significant differences (two-tailed paired Student’s *t-*test, ** *P* < 0.01; * *P* < 0.05; n.s., *P* > 0.05). (**e**) Representative images of inflorescence stems for L*er*, *clv3-1* and *wus-7*. Scale bars = 2 cm. (**f**) Number of siliques per unit stem length of indicated plants. Data represent mean ± SD (*n* = 20). One unit length= 2 cm. Different letters indicate statistically significant differences by one-way ANOVA (Tukey’s multiple comparisons test, *P* < 0.01).

Previous studies revealed that the RM is responsible for the stem elongation ^2,3,36^, we then examined the RMs of all genotypes as previous described ^36^. Compared to WT, the *clv3-1* produced longer RM showing enhanced RM activity, while the *wus-7* generated shorter RM indicative of reduced RM activity (Fig. 1b and c). Previously, we found that the activation of RM was controlled by AP1-TOC1 module, which regulated the circadian stem elogantion^36^. We further investigated the circadian stem elongation of these genotypes. Under continuous light (LL), the *clv3-1* mutant exhibited an increased stem growth rate during the subjective day (ZT0-ZT12 (9:00–21:00)), whereas its growth rate was comparable to that of the WT during the subjective night (ZT12-ZT24 (21:00–9:00)). In contrast, the *wus-7* mutant displayed markedly reduced growth rates throughout the entire diurnal cycle (Fig. 1d). These results indicated that the *CLV3*-*WUS* module is required for both the magnitude and rhythmic pattern of stem elongation. Meanwhile, consistent with their roles in meristem maintenance and organogenesis^45,47^, we found that *clv3-1* developed a larger SAM and produced more siliques relative to WT, whereas *wus-7* formed a smaller SAM and fewer siliques (Fig. 1e, Extended Data Fig. 1d and 1e). Together with the observed defects in circadian-regulated stem elongation, *clv3-1* displayed an elevated number of siliques per unit length compared with WT, while *wus-7* exhibited the opposite phenotype (Fig. 1e and 1f). These results indicate that *CLV3* and *WUS* govern organogenesis and the rhythmic stem elongation by modulating SAM activity to ensure normal shoot morphogenesis.

### TOC1 directly regulate the circadian expression of *CLV3* and subsequent shoot morphogenesis

To understand the underlying mechanisms of *CLV3*-and *WUS*-regulated circadian stem elongation, we examined the expression rhythm of *WUS* and *CLV3* using the IMs of L*er*, sampled as previous described ^36^. RT-qPCR analysis revealed that *WUS* showed an expression rhythm with near to 12-h period in LL conditions, while *CLV3* displayed near 24-h circadian rhythmicity, resembling the circadian pattern of *TOC1* (Fig. 2a and Extended Data Fig. 2a). To determine whether TOC1 regulates the circadian expression of *CLV3*, we assessed *CLV3* rhythmicity in the IMs of the *toc1-17*. Notably, circadian oscillation of *CLV3* was destructed in *toc1-17* (Fig. 2a), suggesting that *TOC1* was crucial for maintaining *CLV3* rhythmicity. Previously, we performed RNA-seq and ChIP-seq analyses and revealed that TOC1 played dual roles in flower development: activating the expression of genes involved in organ development and repressing the expression of genes related to environmental stimulus responses^36^. We then checked the *CLV3* locus and found that two binding peaks of TOC1 located at both upstream and downstream of *CLV3* coding region (Fig. 2b). TOC1 generally bind to the G-box or EE of targets^29,32,36^. Bioinformatic analysis identified several EEs and one G-box located within both the promoter region and 3’-untranslated region (UTR) of *CLV3* (Fig. 2c). ChIP-qPCR assay using the inflorescences of *pTOC1::TOC1-HTF* confirmed that TOC1 directly bound to those EE-containing regions but not the G-box region within the promoter and 3’-UTR of *CLV3* (Fig. 2c and 2d). TOC1 is generally considered as one transcriptional repressor in seedlings^32^. We sought to examine its regulatory effect on *CLV3* using a *LUC* reporter assay^36^. The *pCLV3::LUC::3’UTR* construct was co-transformed with *p35S::TOC1* or *p35S::TOC1-GR* into *Nicotiana benthamiana* leaves (Fig. 2e). Interestingly, TOC1 significantly enhanced LUC activity, and upon TOC1-GR activation, the LUC activity was dramatically induced (Fig. 2e and 2f), indicating that TOC1 directly activated *CLV3* expression. To further determine whether the EEs bound by TOC1 mediate TOC1-dependent activation, we generated site specific mutations into these EEs within the LUC reporter system (Extended Data Fig. 2b). Mutations of EEs in either the promoter or 3’-UTR regions attenuated TOC1-mediated activation of *LUC*, while simultaneous mutations in both regions completely abolished the ability of TOC1 to activate *LUC* expression (Fig. 2g). These findings demonstrated that TOC1 directly activated *CLV3* expression, thereby contributing to the maintenance of the circadian rhythm of *CLV3*.

**Fig. 2.**
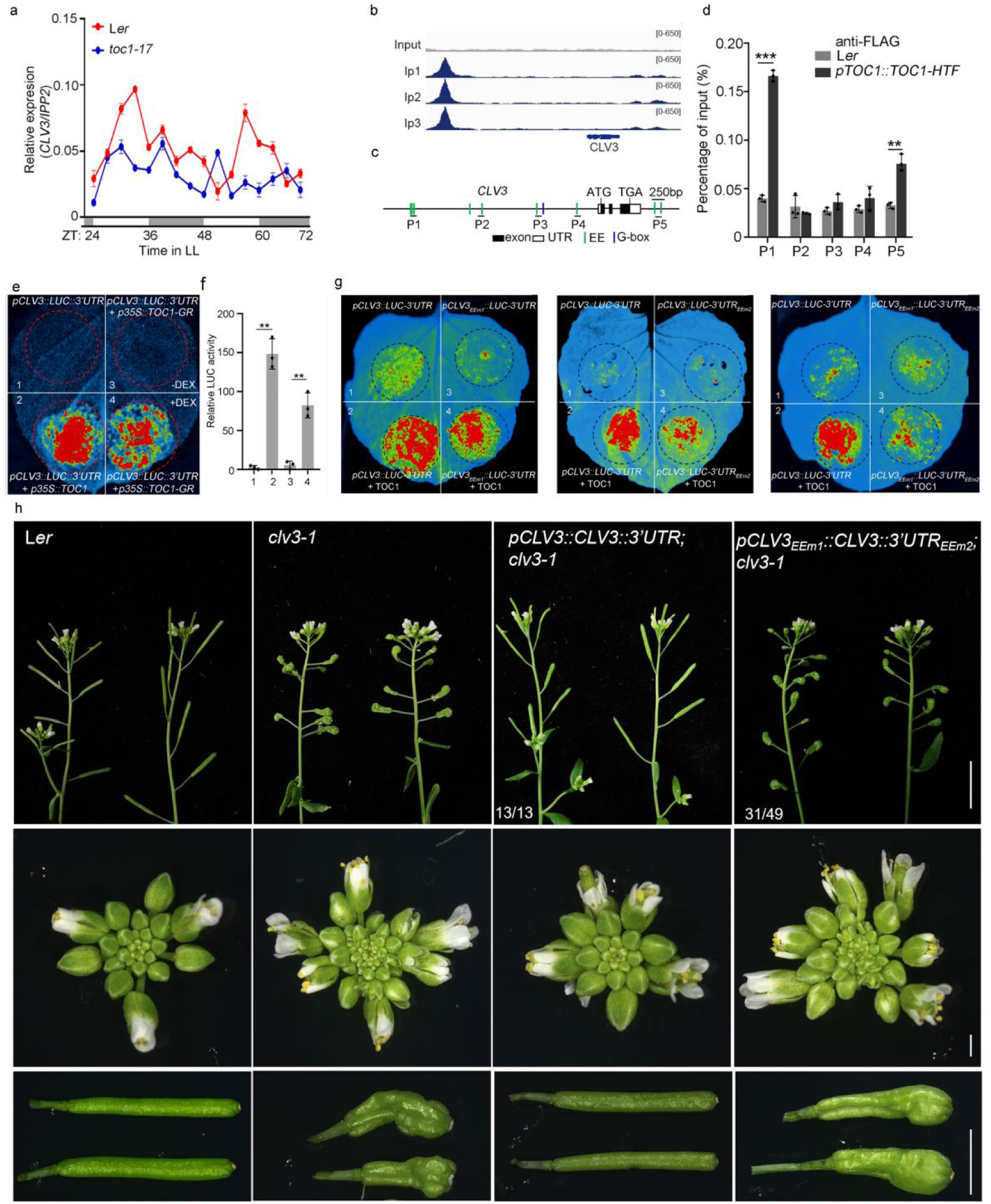
TOC1 directly activates the circadian expression of CLV3. **(a)** RT-qPCR to examine the circadian expression patterns of *CLV3* in meristems of L*er* and *toc1-17*. Data represent mean ± SD of three biological replicates. *IPP2* was used as a normalization control. White and gray bars represent subjective day and night, respectively. (**b**) ChIP-seq analysis revealed a significant binding peak of TOC1 in the upstream region of the *CLV3* promoter and another binding peak in the 3′UTR region (the top panel shows the input, and the bottom panel shows three biological replicates of the IP). **(c)** Diagram of the *CLV3* locus. White and black rectangles and black lines represent untranslated regions, coding regions, and introns, respectively. Green lines and blue line represent EE motifs and G-box, respectively. The regions examined for TOC1 binding are indicated (P1-P5). **(d)** ChIP-qPCR to examine the binding of TOC1 to *CLV3* in the meristems of indicated plants. ChIP signal was quantified as the percentage of total input DNA by qPCR. The examined loci were indicated in (c). Data represent mean ± SD from three biological replicates. Student’s *t*-test; \*\*\**P* < 0.001, \*\**P* < 0.01. **(e-f)** Luciferase complementation imaging (LCI) system (e), and quantitative analysis of LUC activity (f), to examine the direct activation of *TOC1* on *CLV3* expression. The *pCLV3::LUC-3’UTR* construct was co-transformed into tobacco leaves either without (1) or with (2) *p35S::TOC1*. In parallel, *pCLV3::LUC-3’UTR* was co-transformed with *p35S::TOC1-GR*, followed by DMSO (3) or DEX (4) treatment. For quantitative analysis, the LUC activity of *pCLV3::LUC-3’UTR* transformed alone was set as 1. Three independent experiments were conducted with similar results. Student’s *t*-test (two-tailed); \*\**P* < 0.01. **(g)** Luciferase complementation imaging (LCI) assay the direct activation of *TOC1* on *CLV3* expression. The *pCLV3::LUC-3’UTR* construct was co-transformed into tobacco leaves without (1) or with (2) *p35S::TOC1*. The *pCLV3_EEm1_::LUC-3’UTR*, *pCLV3::LUC-3’UTR_EEm2_*, or *pCLV3_EEm1_::LUC-3’UTR_EEm2_* constructs were co-transformed into tobacco leaves without (3) or with (4) *p35S::TOC1*. Three independent experiments were conducted with similar results. **(h)** Phenotypes of whole plants, inflorescences, and siliques of indicated plants. Scale bars: 1.5 cm.

We further investigated how TOC1 modulates *CLV3* function. Expression of *CLV3* driven by its native promoter and 3′-UTR (*pCLV3::CLV3::3′UTR*) fully rescued the developmental defects of the *clv3-1* mutant, including impaired shoot morphogenesis caused by an enlarged SAM, excess silique and carpel production, and moderately elongated stems (Fig. 2h and Extended Data Fig. 2c-g). By contrast, the phenotypes of most *pCLV3_EEm1_::CLV3::3′UTR_EEm2;_clv3-1* transgenic lines remained similar to that of the *clv3-1* mutant (Fig. 2h and Extended Data Fig. 2c-g). Collectively, these results indicate that TOC1-regulated rhythmic *CLV3* expression is essential for its roles in controlling meristem activity and stem elongation. Previously, we discovered that *TOC1* is required for the RM activation and stem elongation^36^. Compared with WT, *clv3-1;toc1-17* produced moderately elongated stem similar to *clv3-1*, but instead of the dwarf *toc1-17*, indicating that *CLV3* acts downstream of *TOC1* to regulate stem elongation (Extended Data Fig. 2h). To assess the effect of rhythmic *CLV3* expression on the role of *TOC1* in stem elongation, we treated the inflorescence of *toc1-17* with a synthesized CLV3 peptide, known to function in SAM maintenance ^48,49^, with different patterns. Daily CLV3 treatment applied at ZT12 (21:00), shortly after its expression peak (Fig. 2a), partially rescued the stem-elongation defect in *toc1-17* mutants. By contrast, daily CLV3 treatment at ZT0 (9:00) or 6-hour periodic CLV3 treatments (ZT 0/6/12/18 (9:00/15:00/21:00/3:00)) failed to restore this defect (Extended Data Fig. 2i). These results indicate that CLV3 is required for TOC1-mediated regulation of stem elongation.

### *TOC1*-rhythmic *CLV3* expression is sufficient for organogenesis and stem elongation

To investigate the role of circadian rhythmicity of *CLV3* in meristem maintenance and stem elongation, we ectopically expressed *CLV3* driven by the promoter of *TOC1* or *CCA1* in *clv3-1*, thereby mimicking *TOC1*-like or *CCA1*-like circadian expression patterns. Remarkably, transgene *pTOC1::CLV3* nearly fully rescued the developmental defects of *clv3-1*, restoring normal inflorescence structure, carpel number, and RM size to levels comparable to WT. In contrast, *pCCA1::CLV3;clv3-1* displayed markedly delayed flowering time with enlarged inflorescence, increased carpel number and decreased RM size, and a complete loss of stem development compared to both WT and *clv3-1* (Fig. 3a 3b, and Extended Data Fig. 3a). In addition, crossing-mediated transduction of *pTOC1::CLV3* into the *toc1-17* mutant rescued its stem-elongation defect, consistent with observations from exogenous CLV3 treatment of *toc1-17* (Extended Data Fig. 3b-d). Analysis of expression patterns confirmed that the circadian expressions of *CLV3* in the respective transgenic lines were similar to the circadian rhythm of *TOC1* and *CCA1*, respectively (Fig. 3c and Extended Data Fig. 3e). These findings suggested that the circadian expression pattern of *CLV3* was required for its role in regulating meristem activity and promoting stem elongation.

**Fig. 3.**
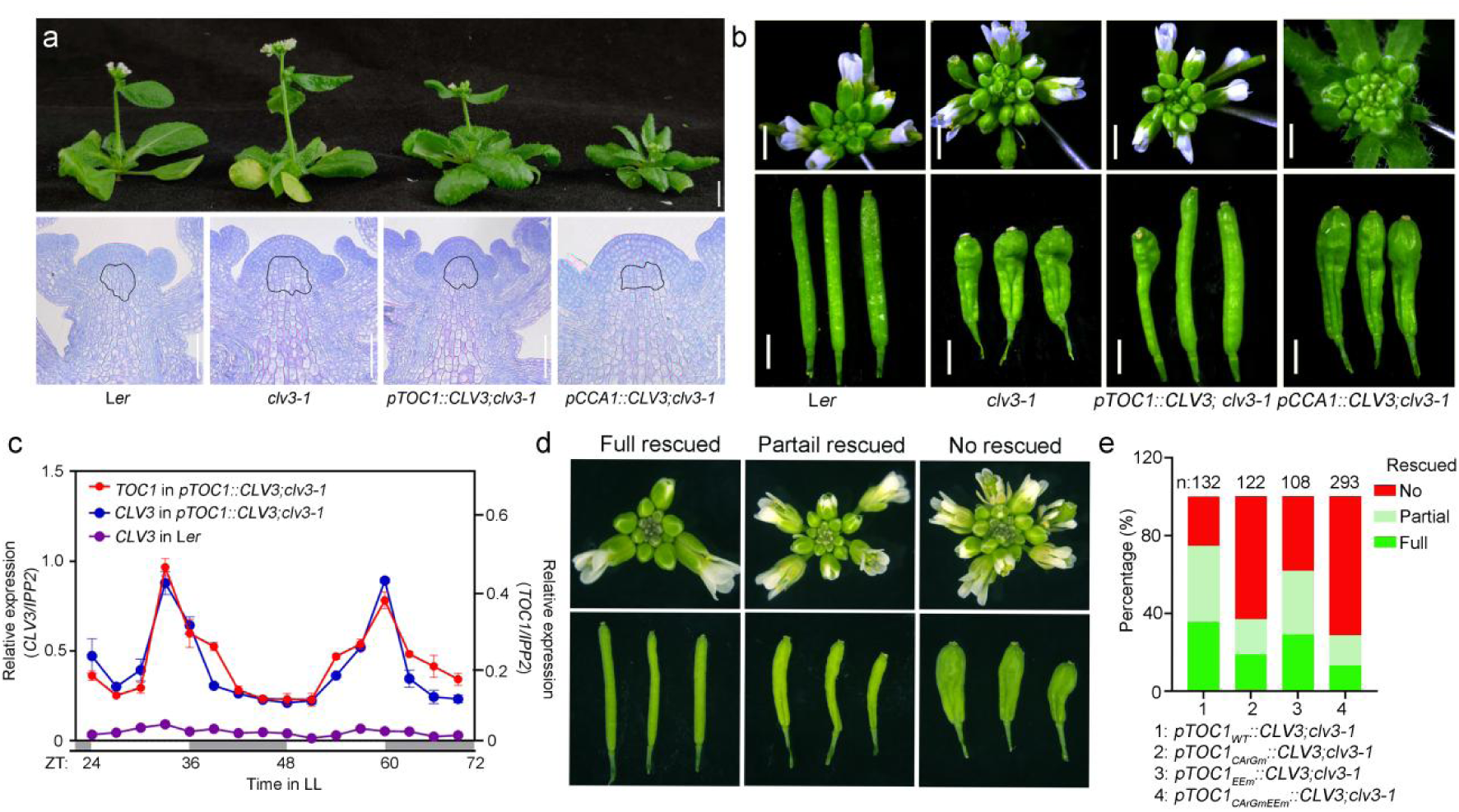
Rhythmic *CLV3* expression is sufficient for organogenesis and stem elongation. **(a)** Phenotypes of plants and longitudinal sections of shoot apices of indicated plants showing the range of RM. The black line outlines the RM region. Representative section from 15 samples per genotype is shown. Scale bars for whole plants = 1cm. Scale bar for longitudinal shoot-apex sections = 100 μm. **(b)** Phenotypes of inflorescences and siliques of indicated plants. Scale bars: 1 cm. **(c)** RT-qPCR to examine the expression patterns of *TOC1* in *pTOC1::CLV3;clv3-1* (left y-axis), and *CLV3* in L*er* and *pTOC1::CLV3;clv3-1* (right y-axis). Inflorescences were sampled at 3-hour intervals as shown in (c). Data represent mean ± SD of three biological replicates. *IPP2* was used as a normalization control. White and gray bars represent subjective day and night, respectively. **(d)** Inflorescence and siliques phenotypes of indicated plants. Phenotype was classified into three groups based on the number of carpels: fully rescued (2 carpels/siliques), partially rescued (3-4 carpels/siliques) with normal inflorescence, and non-rescued lines with enlarged inflorescence and more than 4 carpels/siliques. **(e)** Percentage of plants exhibiting distinct phenotypes shown in (d) and *n* values for each genotype are labelled on top of each bar.

We previously demonstrated that AP1 physically interacts with CCA1 to sustain *TOC1* circadian expression via CArG-box and EE motifs within its promoter, and TOC1 non-cell-autonomously modulates rhythmic stem elongation ^36^. To further assess whether TOC1-controled circadian expression of *CLV3* mediate AP1/CCA1-TOC1 regulatory circuit in meristem maintenance, we transformed *clv3-1* with *pTOC1_WT_::CLV3, pTOC1_CArGm_::CLV3, pTOC1_EEm_::CLV3* and *pTOC1_CArGm EEm_::CLV3,* respectively, each containing site-directed mutations in the CArG-box or/and EEs within the *TOC1* promoter (Extended Data Fig. 3f). The resulting transgenic plants were classified into three phenotypic groups based on the number of carpels: fully rescued (2 carpels/silique), partially rescued (3-4 carpels/silique) with normal inflorescence, and non-rescued lines with enlarged inflorescence and more than 4 carpels/silique (Fig. 3d). While approximately 80% of *pTOC1_WT_::CLV3;clv3-1* displayed either fully or partially rescued phenotype, the introduction of single site mutations at the CArG-box or EEs significantly reduced the percentage of rescued lines. Moreover, simultaneous mutations at both regulatory elements dramatically increased the percentage of transgenic plants with non-rescued phenotype (Fig. 3e). These results indicated that AP1-CCA1 controlled circadian regulation of *TOC1* which was essential for the function of *CLV3* on meristem activity maintenance. To validate the dependency of TOC1-regulated *CLV3* activity on AP1 and CCA1, we crossed a fully rescued *pTOC1_WT_::CLV3;clv3-1* line into *cca1-8* and *ap1;cal*, respectively, generating *pTOC1_WT_::CLV3;clv3-1;cca1-8* and *pTOC1_WT_::CLV3;clv3-1;ap1;cal* lines: Both lines displayed a reversion to the *clv3-1* phenotype, as evidenced by showing enlarged inflorescence and increased carpel numbers (Extended Data Fig. 3g). These findings demonstrated that the functional output of TOC1-regulated *CLV3* depended on both *AP1* and *CCA1*.

### Rhythmic expression pattern rather than transcript abundance of *CLV3* governs

#### *WUS* expression and SAM activity maintenance

To investigate how circadian expression of *CLV3* influences meristem maintenance and stem growth, we first constitutively expressed *CLV3* under the control of CaM35S promoter in WT. In all transgenic plants analyzed (*n*=23), the SAMs were arrested or terminated after the initiation of several true leaves, phenocopying the *wus* mutant phenotype, as previously reported (Extended Data Fig. 4)^46,47,50^. RT-qPCR analysis revealed that *CLV3* transcript levels in *p35S::CLV3* were nearly 100-fold higher than in WT, accompanied by a drastic reduction in *WUS* expression. These findings were corroborated by *in situ* hybridization (Fig. 4a and 4b), demonstrating that constitutive and spatially uniform expression of *CLV3* repressed *WUS* expression, ultimately leading to the termination of stem cell. Unexpectedly, we noticed a substantial increase in *CLV3* expression in both *pTOC1::CLV3;clv3-1* and *pCCA1::CLV3;clv3-1* lines, which displayed either rescued or enhanced *clv3-1* phenotypes, respectively, rather than *wus* phenotype (Fig. 3a). In the WT, *WUS* and *CLV3* were spatially restricted and normally expressed in the OC and CZ, respectively^51^. Nevertheless, in *clv3-1*, the disruption of *CLV3*-*WUS* negative feedback loop resulted in expanded expression domains and increased transcript levels of both genes due to compensatory mechanisms^51,52^. In *pTOC1::CLV3;clv3-1*, the spatial expression domain and transcript level of *WUS* were restored to patterns resembling those in WT (Fig. 4b), correlating with the rescued phenotype (Fig. 3a). Notably, despite this restoration, *CLV3* expression remained elevated in both domain size and intensity, indicating that both the spatial extent and strength of *CLV3* expression were still enhanced. In contrast, *pCCA1::CLV3;clv3-1* showed significantly expanded and intensified expression domains of both *CLV3* and *WUS*, particularly in IMs and FMs, as confirmed by RT-qPCR analysis (Fig. 4a and 4b). These results indicated that the correlation between expressions of *CLV3* and *WUS* expression was disrupted, implying that the circadian expression pattern of *CLV3* played a more crucial role than its expression intensity in maintaining the *CLV3*-*WUS* regulatory feedback loop.

**Fig. 4.**
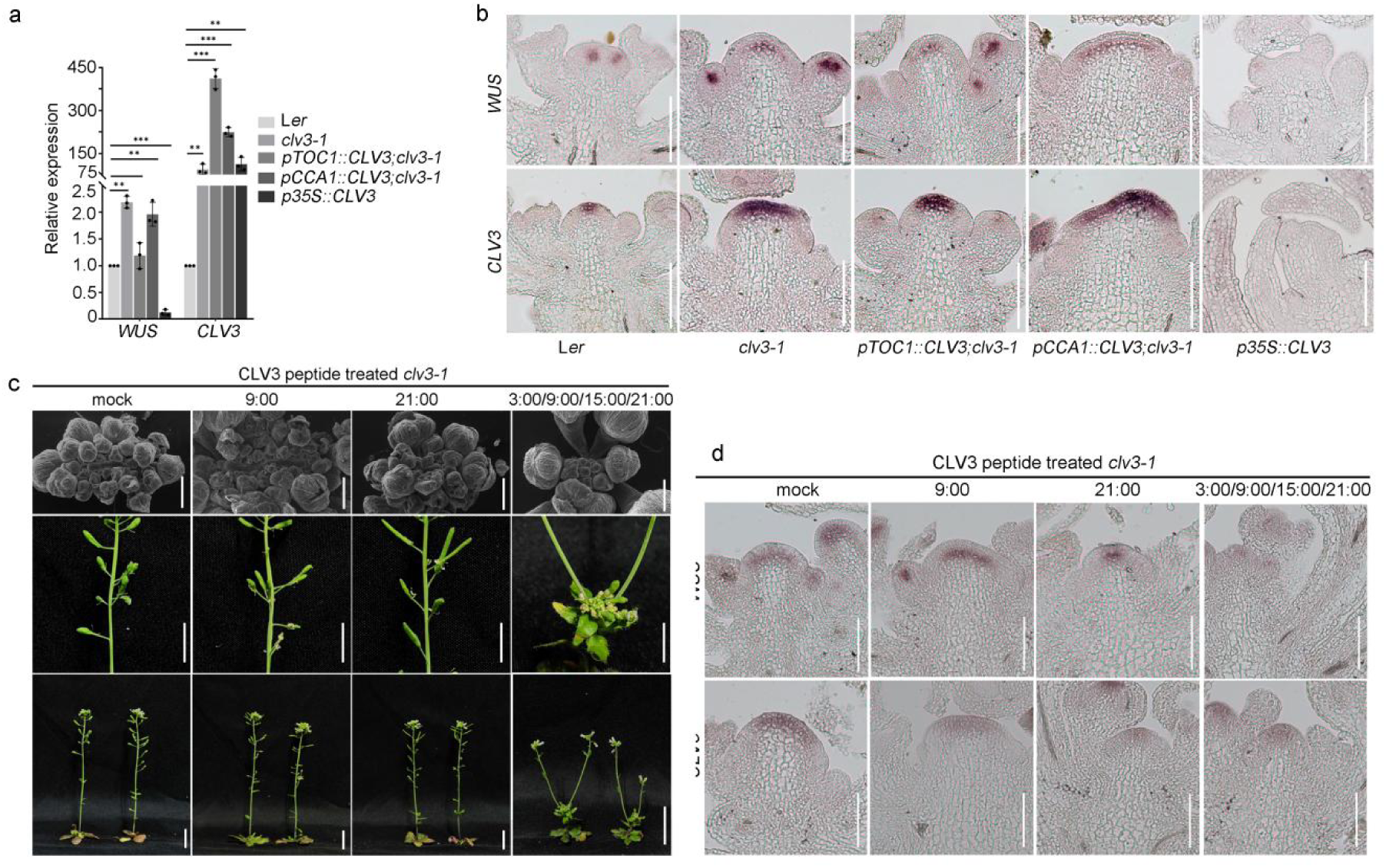
Circadian expression of *CLV3* is crucial for the CLV3-WUS regulatory loop. **(a)** RT-qPCR to examine the expression level of *WUS* and *CLV3* in indicated plants. Gene expression in L*er* was set as 1. Data represent mean ± SD of three biological replicates. *IPP2* was used as a normalization control. Student’s *t*-test (two-tailed); \*\*\**P* < 0.001; \*\**P* < 0.01. **(b)** *In situ* hybridization to detect expression patterns of *WUS* and *CLV3* in meristems of indicated plants. Scale bars: 100 μm. **(c)** IMs, Inflorescences, siliques and whole plants phenotypes of CLV3 peptide treated *clv3-1* at indicated time points for 7 days. The plants grown under 12/12(light/dark) conditions for additional 10 days after treatment followed by photograph. Scale bars: 0.5 cm. **(d)** *In-situ* hybridization to detect expression patterns of *WUS* and *CLV3* in IMs of *clv3-1* from (c). Scale bars: 100 μm.

To further investigate whether the circadian accumulation of *CLV3* peptide regulates meristem maintenance and activity, we treated the SAMs of *clv3-1* with CLV3 peptide at different time points for 7 days. While constitutive treatment with DMSO (mock treatment) had no effect on the SAM development of *clv3-1*, periodic treatment with CLV3 every 6 hours (ZT 0/6/12/18 (9:00/15:00/21:00/3:00)) resulted in SAM termination, consistent with previous findings ^48^. Strikingly, CLV3 application at ZT12(21:00) rescued the developmental defects of *clv3-1*, restoring normal carpel number and inflorescence size, and reducing SAM size to levels comparable to the WT. In contrast, treatment at ZT0 (9:00), when *CLV3* was expressed at its minimum level, resulted in the enhanced phenotype of *clv3-1* resembling to that of *pCCA1::CLV3;clv3-1* plants, with increased carpel and silique numbers and enlarged inflorescence size with fasciate SAM (Fig. 4c). *In situ* hybridization analysis revealed that constitutive treatment of CLV3 repressed the expression of both *WUS* and *CLV3*, leading to SAM termination. Conversely, CLV3 treatment at ZT12 (21:00) restored in *clv3-1* expression domains and intensities of *CLV3* and *WUS* to levels similar to those in WT, along with a normalized SAM size. In contrast, CLV3 treatment at ZT0 (9:00) led to increased *WUS* expression and decreased *CLV3* expression (Fig. 4d). These results suggested that the circadian timing of *CLV3* accumulation was critical for its functions within the *CLV3*-*WUS* regulatory circuit and meristem maintenance. Moreover, the data implies the presence of a rapid turnover or degradation mechanism for CLV3 peptide in meristems.

#### CLV3 temporally regulates rhythmic accumulation of WUS

These findings prompted us to examine whether WUS protein also exhibits rhythmic accumulation in the meristems. In contrast to the ∼12-h oscillatory expression pattern in the WT, *WUS* expression becomes arrhythmic in the *clv3-1* mutant (Extended Data Fig. 5a). Due to the restricted expression of *WUS* in the OC, assessing its circadian expression accurately using RT-qPCR was challenging and unreliable. To overcome this limitation, we examined rhythmic WUS accumulation by quantitating GFP fluorescence signals in *pWUS::WUS-GFP;wus-101* ^53^. The expression domain and protein accumulation of WUS-GFP reached the maximum level at ZT0 (9:00), then gradually and slowly decreased to a minimum level at ZT12 (21:00), and subsequently increased rapidly to the maximum level again at the next ZT20 until ZT24, displaying a near 24-hour circadian oscillation pattern, which was confirmed by Western blotting assay using anti-GFP antibody to examine WUS-GFP accumulation (Fig. 5a-5c). Together with the approximately 12-hour rhythmic expression pattern of *WUS* (Extended Data Fig. 2a), the results suggested that *WUS* was subjected to different regulatory mechanisms orchestrating its transcription and protein accumulation, consistent with previous findings ^16^. We then examined the regulation of CLV3 on *WUS* expression and WUS accumulation. Single CLV3 treatment at ZT12(21:00) resulted in rapid repression of *WUS* expression that was restored after 12-hour treatment, while single CLV3 treatment at ZT0(9:00) led to a stable but mild repression of *WUS* expression (Extended Data Fig. 5b). Meanwhile, the WUS-GFP accumulation was unchanged at both daytime and nighttime in the *pWUS::WUS-GFP;wus-101;clv3-1*, which was confirmed by Western Blotting assay (Fig. 5a, 5b and 5d). These results indicated that *CLV3* dynamically regulates *WUS* expression and is required for the inhibition of WUS accumulation at daytime.

**Fig. 5.**
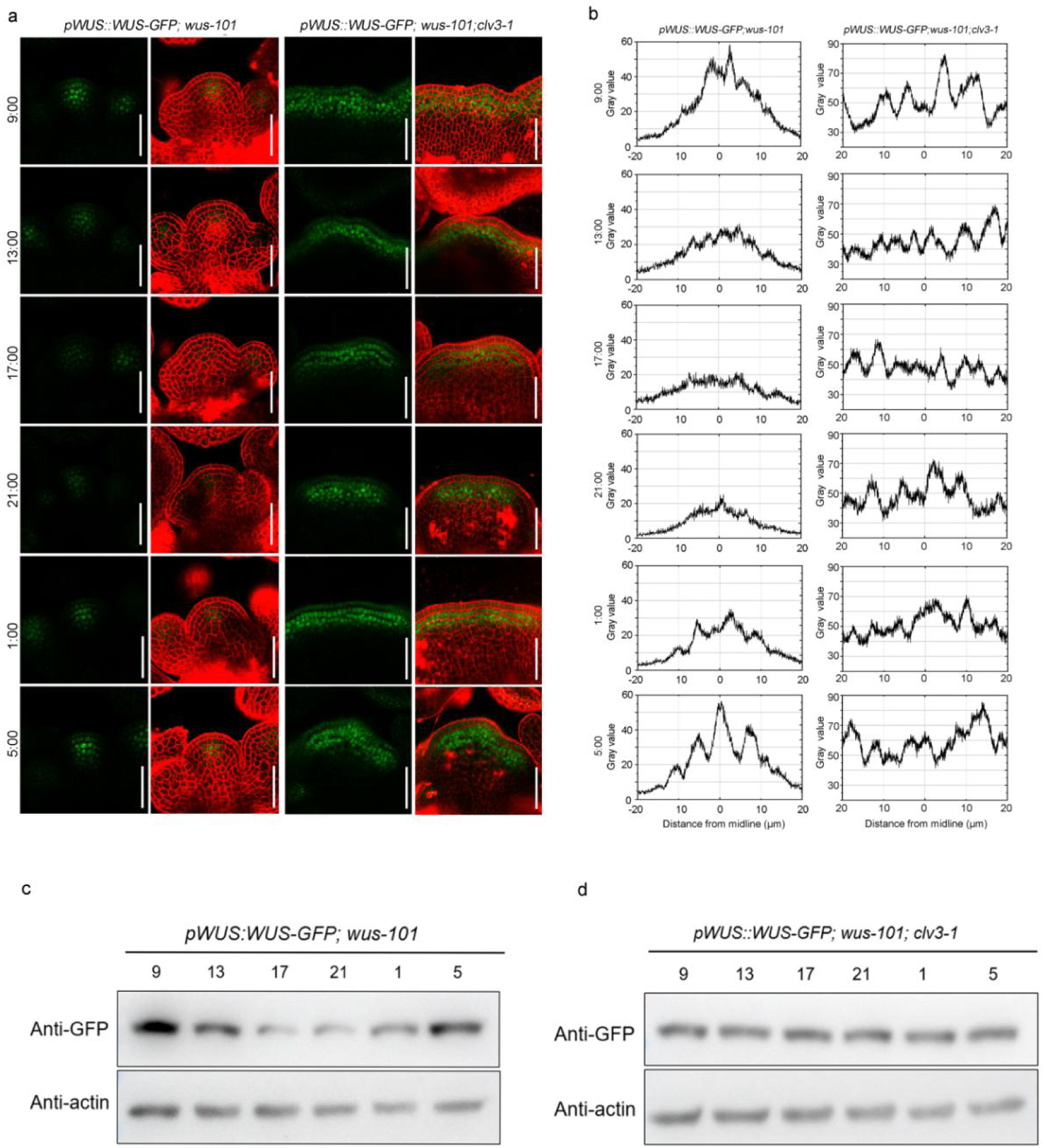
CLV3 temporally regulates the rhythmic accumulation of WUS. **(a)** WUS-GFP fluorescence accumulation in IMs of *pWUS::WUS-GFP; wus-101* and *pWUS::WUS-GFP; wus-101; clv3-1*. The plants were grown under 12/12 (light/dark) conditions, and IMs from plants at similar developmental stages were collected for GFP examination at indicated time points (*n* = 10). Scale bars = 50 μm. **(b)** Quantitative analysis of WUS-GFP fluorescence intensity in IMs of indicated plants at the indicated time points shown in (a). **(c-d**) Meristem samples were collected every 4 h over a 24 h period in LL from the plants in (a). Total proteins were separated by SDS-PAGE, and WUS-GFP was detected using an anti-GFP antibody. Anti-actin antibody was used as the loading control. Three independent experiments were conducted with similar results.

#### WUS acts downstream of AP1/CCA1-TOC1-CLV3 cascade in stem elongation

Previous studies have demonstrated that increased *WUS* level induces the expansion of the CZ and the cell division rate of cells in PZ, whereas reduced *WUS* levels promotes cell differentiation in the PZ and possibly in the RM^10^. To further investigate the role of *WUS* in circadian stem growth, we conducted genetic analyses. The *wus-7* mutant showed reduced stem, a loss of circadian growth rhythms, and a reduced RM size compared to WT, and transgene of *pTOC1::CLV3*, *pTOC1::TOC1-GFP* and *pAP1::TOC1-GFP* were able to rescue the developmental defects of *clv3-1* and *toc1-17* mutants, respectively (Fig. 1a-1c and Fig. 6a-6c)^36^. The corresponding triple mutant lines: *pTOC1::CLV3;clv3-1;wus-7*, *pTOC1::TOC1-GFP;toc1-17;wus-7*and *pAP1::TOC1-GFP;toc1-17;wus-7* all phenocopied the phenotype of *wus-7* mutant (Fig. 6a-6c). Moreover, rhythmic stem elongation was abolished in all genotypes under the wus mutant background (Fig. 6d). Collectively, these findings demonstrated that *WUS* worked as a central downstream effector of *AP1*, *TOC1* and *CLV3* in regulating circadian stem development.

**Fig. 6.**
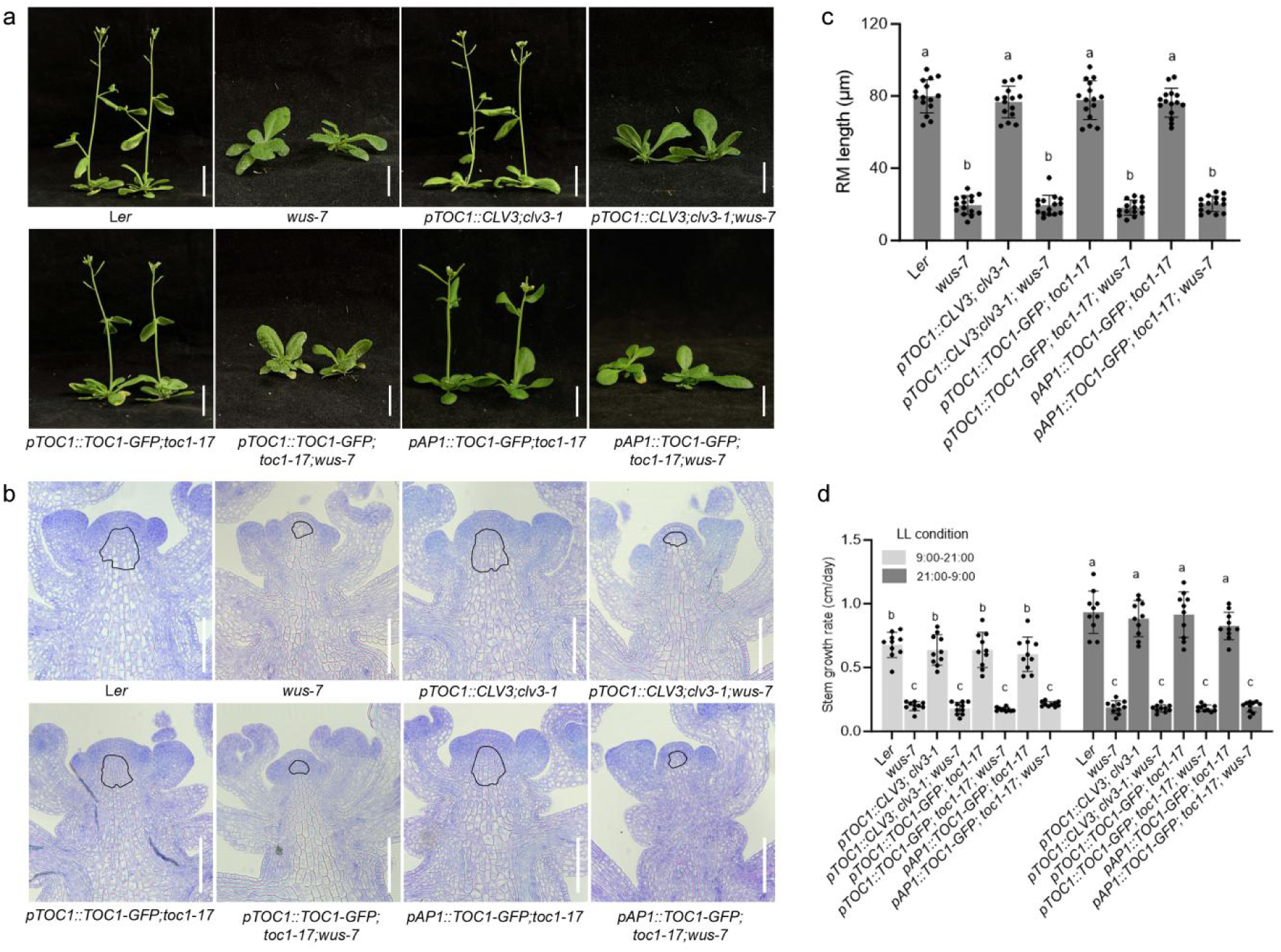
WUS acts downstream of the AP1/CCA1-TOC1-CLV3 cascade in stem elongation. **(a)** Phenotypes of indicated plants. Scale bars: 1 cm. **(b)** Phenotypes of longitudinal sections of shoot apices of indicated plants showing the range of RM. The black line outlines the RM region. Representative section from 15 samples per genotype is shown. Scale bars: 100 μm. **(c)** Statistical analysis of the length of RM of indicated plants. Data represent mean ± SD (*n* = 15). Different letters indicate statistically significant differences by one-way ANOVA (Tukey’s multiple comparisons test, *P* < 0.01). **(d)** Stem growth rate of plants grown under LL conditions of indicated plants. 9:00-21:00 subjective day; 21:00-9:00 subjective night. The letters indicate the significance groups at *P* < 0.05 (one-way ANOVA and Tukey’s test). Different letters between genotypes and day/night conditions indicate significance.

#### WUS regulates cell wall properties through controlling PMEs expression

Previous biochemical and transcriptome analyses revealed that WUS acts as either transcriptional repressor or activator to maintain SAM activity by controlling genes involved in cell division, auxin signaling and meristem-related processes^54,55^. In particular, RNA-seq and ChIP-seq data from *35S::WUS-GR* plants indicated that WUS broadly modulates auxin pathway activities, from transport and perception to signal transduction and transcriptional responses ^56^. To gain deeper insights into the molecular mechanism by which WUS regulates SAM development, we systematically reanalyzed previously reported omics datasets ^56^. RNA-seq analysis identified hundreds of differentially expressed genes (Extended Data Table 1.1). Gene Ontology (GO) enrichment revealed that most of these genes are involved in multiple responses to diverse stimuli, including oxygen availability, cold and heat stress, light signaling, and sugar metabolism, as well as leaf developmental processes (Extended Data Fig. 6a and 6b). Although the analysis of omics data was primarily derived from seedlings with ectopic *WUS* expression, which makes it difficult to determine whether the regulated biological pathways occur *in vivo* ^56^, we still found that WUS-repressed genes are predominantly enriched in cell wall developmental processes, including cell wall thickening and callose deposition, indicating that WUS modulates cell wall properties by suppressing cell wall rigidity. Therefore, we focused on the identified target loci that were bound by WUS (Extended Data Table 1.2) ^56^. GO analysis showed that the WUS-binding loci were highly involved in multiple developmental processes, such as meristem and floral development, response to auxin and cell division, as well as cell wall organization and biogenesis (Extended Data Fig. 6c).

Given that PMEs were critical for the SAM organization and WUS-controlled PMEs is essential for sustaining stem cell identity and SAM organization^43,57^, we isolated several *PMEs* that were bound by WUS directly (Extended Data Fig. 6d), indicating that WUS regulates cell wall properties by controlling expression of *PMEs*. One candidate target gene, PME5, was further investigated ^57^. Dramatic enrichment of WUS was found at the promoter of *PME5* containing one G-box element that was bound by WUS, which was verified by ChIP-qPCR (Fig. 7a-7c)^57,58^. We then constructed *pPME5::LUC* and *pPME^5G-boxm^::LUC* reporter constructs harboring point mutations in the G-box motif. These reporters were individually co-transformed with *p35S::WUS-GFP* into *Nicotiana benthamiana* leaves. We observed that WUS-GFP substantially elevated *LUC* activity in leaves expressing *pPME5::LUC*, whereas the G-box mutation markedly attenuated WUS-mediated activation of *PME5* expression (Fig. 7d and 7e), demonstrating that WUS could directly induce the expression of *PME5*. Consistently, *PME5* expression was decreased in *wus-7* (Fig. 7f). To investigate the role of *PME5* mediating the function of *WUS* in RM activation and stem development, we overexpressed *PME5* in both WT and *wus-7*. The stem of *PME5-OE* plant was longer than that of WT, in line with previous report ^40^, indicating that *PME5* enhances stem elongation. Intriguingly, *PME5* overexpression could rescue the stem developmental defect of *wus-7* (Fig. 7g). Consistently, the RM of *PME5*-overexpressing plants was longer than that of wild-type plants. Meanwhile, the RM in *PME5-OE wus-7* double mutants was also longer than that of *wus-7* single mutants (Fig. 7g and 7h), indicating that *PME5* promotes stem development by activating RM activity.

**Fig. 7.**
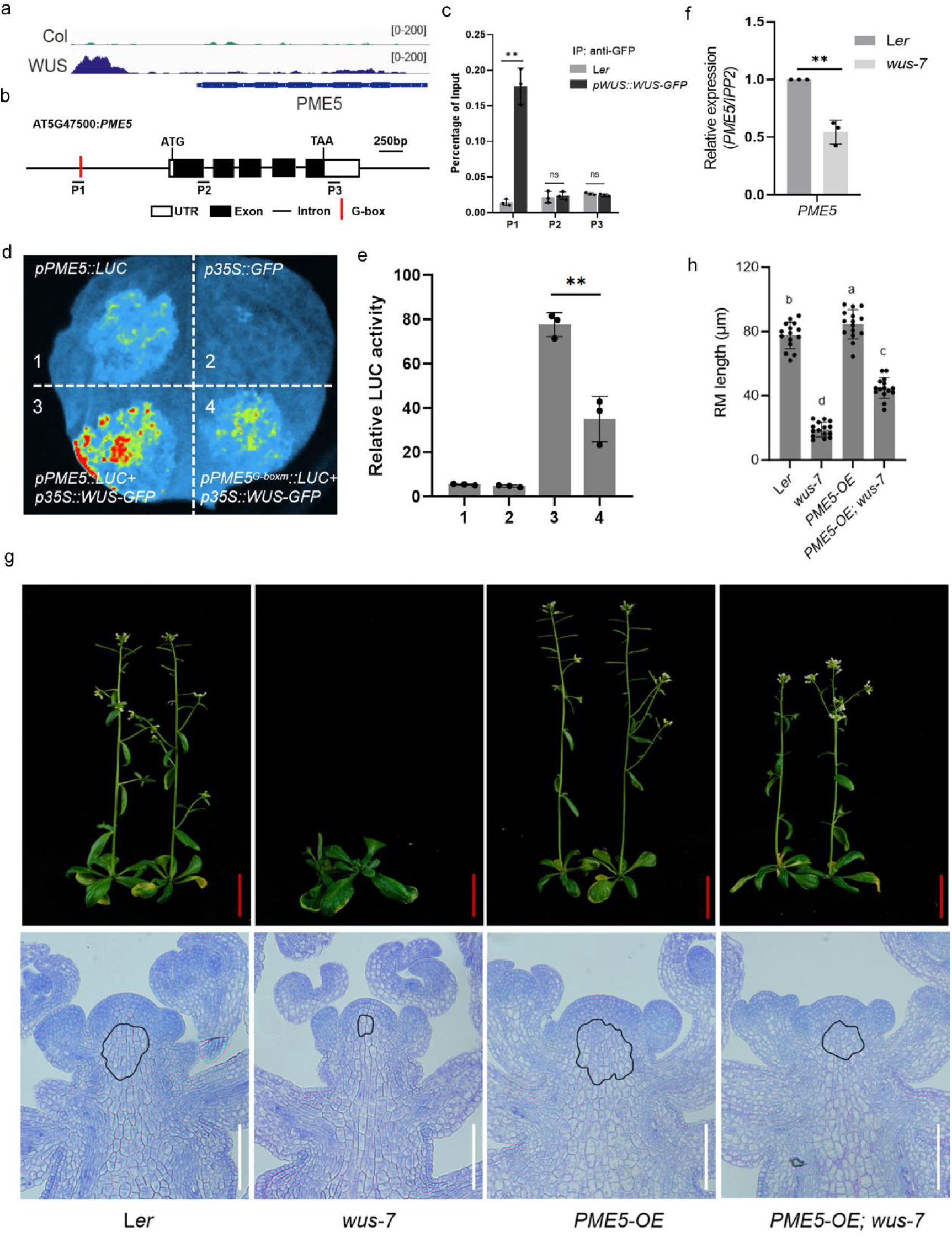
WUS regulates stem elongation through controlling *PME5* expression. **(a)** ChIP-seq analysis revealed a significant binding peak of WUS in the upstream region of the *PME5* promoter region (the top panel shows the input, and the bottom panel shows three biological replicates of the IP). **(b)** Diagram of the *PME5* locus. White and black rectangles and black lines represent untranslated regions, coding regions, and introns, respectively. The red line represents the G-box motif. The regions examined for WUS binding are indicated (P1-P3). **(c)** ChIP-qPCR quantification of WUS enrichment at the *PME5* locus in *pWUS::WUS-GFP* plants. Enrichment values are shown as percentage of input DNA. Plants without the *WUS-GFP* transgene served as the negative control. Data represent mean ± SD of three biological replicates. Two-tailed Student’s *t-*test; \*\**P* < 0.01; n.s., not significant. **(d-e**) LCI system (d), and quantitative analysis of LUC activity (e), to examine the direct activation of WUS on *PME5* expression. The *pPME5::LUC* construct was co-transformed into tobacco leaves without (1) or with (3) *p35S::WUS-GFP*. The *pPME5_G-boxm_::LUC* construct was co-transformed into tobacco leaves together with *p35S::WUS-GFP* (4). *p35S::GFP* alone served as the negative control (2). Three independent experiments were conducted with similar results. Student’s *t*-test (two-tailed); \*\**P* < 0.01. **(f)** RT-qPCR to examine the expression level of *PME5* in L*er* and *wus-7*. Gene expression in L*er* was set as 1. Data represent mean ± SD of three biological replicates. *IPP2* was used as a normalization control. Student’s *t*-test (two-tailed); ** *P* < 0.01. **(g)** Phenotypes of whole plants and longitudinal sections of shoot apices of indicated plants showing the range of RM. The black line outlines the RM region. Representative section from 15 samples per genotype is shown. Red scale bars: 1 cm; white scale bars: 100 μm. **(h)** Statistical analysis of the length of RM of indicated plants. Data represent mean ± SD (*n* = 15). Different letters indicate statistically significant differences by one-way ANOVA (Tukey’s multiple comparisons test, *P* < 0.01).

## Discussion

Development of a vertical shoot axis (stem) is one of the most successfully evolutionary events for modern land plants, which enables plants grow from two-dimensional to three-dimensional pattern to establish optimal shoot architecture ^19^. All post-embryonic above-ground organs are derived from the SAM. The SAM architecture and synchronized floral organogenesis combined with rhythmic stem elongation collectively shape the canonical plant shoot morphology. The spatially restricted *CLV3*-*WUS* negative feedback loop serves as the core regulatory module was well characterized in meristem maintenance and organogenesis ^59^, while the mechanisms underlying the rhythmic stem elongation remains poorly understand. Compared to the studies focused on stem secondary growth and vascular development, relatively little attention has been given to primary stem growth despite the classic morphological and physiological studies of early stem development conducted in the 1950-60s ^2^.

In *Arabidopsis thaliana*, the RM remains quiescent during the vegetative stage and is activated to initiate stem development during floral transition ^3^. Recently, our work revealed that AP1, one of FM and floral organ identity genes, physically interacts with the morning oscillator CCA1 to dynamically modulate *TOC1* circadian transcription in FMs. TOC1 non-cell-autonomously controls stem elongation partially through regulating *VGD1* expression in IMs ^36^. Whether and how *TOC1* orchestrates organogenesis and rhythmic stem growth remain unclear. Classical models established that the feedback regulatory *CLV3*-*WUS* loop maintains SAM size and orchestrates floral primordia initiation at the peripheral zone^59^. Our work here further demonstrates that this conserved spatial circuit simultaneously controls RM activity, which drives post-floral transition stem elongation, thereby coupling organ formation and stem elongation to construct intact aerial plant architecture. The *clv3-1* allele eliminates CLV3-mediated suppression of WUS, leading to enlarged SAMs with excessive floral organ production and hyperactive RM cells that generate elongated inflorescence stems. In contrast, the *wus-7* mutant exhibits compromised stem cell pools, suppressed organogenesis and severely shortened stems (Fig. 1), highlighting that balanced *CLV3*-*WUS* signaling is indispensable for matching organ number and stem growth magnitude. These results suggested that SAM activity homeostasis are critical for optimal shoot architecture formation.

The role of circadian clocks in adult stem cell maintenance and tissue regeneration has been widely studied. In mammals, the circadian rhythms are orchestrated by a hierarchical network of oscillators, with the suprachiasmatic nucleus acting as the central pacemaker coordinating a number of independent oscillators ^60^. Similarly, in plants, the shoot apex has been proposed to act analogously to the suprachiasmatic nucleus, influencing root circadian activity ^27^. Despite these advances, the mechanisms by which the circadian clock regulates meristem activity remains largely enigmatic. We discovered here that TOC1 directly activated *CLV3* expression and maintained the circadian expression pattern of *CLV3*, which is critical for its role in normal meristem activity and organogenesis (Fig. 2 and 3). Moreover, point mutations in EEs and CArG-box within the *TOC1* promoter, binding sites for CCA1 and AP1, impaired the rescue effect of *pTOC1::CLV3* (Fig. 3d-3e), demonstrating that the functions of CLV3 rhythmicity were controlled by AP1-CCA1 module via TOC1. While TOC1 is widely known as a transcriptional repressor in the vegetative stage ^29,32^, our recent transcriptomic analyses show it acts as both an activator and repressor during flower development. TOC1 also directly activates *VGD1* to drive stem elongation ^36^. Integrating our CLV3 data and previously reported rice *OsTGAL3a/b* ^35^, these observations indicate that TOC1 may switch between repressive and activating roles, probably through association with different co-regulators. Whether TOC1-CLV3 module and its downstream oscillators relay the effects of SAM on the other organs is an attractive question waiting for further investigation.

Spatially feedback regulatory loop of *CLV3*-*WUS* between OC and CZ is important for the meristem maintenance. Meanwhile, both the nuclear levels and the spatial distribution of WUS dynamically controlled by CLV3, are critical for maintaining CZ identity and meristem proliferation regulation^14,16,61,62^. However, the temporal regulation of *CLV3*-*WUS* loop has received comparatively less attention. Beyond static spatial homeostasis, we revealed here that the circadian clock integrates timing cues into the *CLV3*-*WUS* circuit via the FM-derived AP1/CCA1-TOC1 cascade, establishing rhythmic growth patterns. Our data demonstrate that the rhythmic expression pattern of *CLV3*, rather than its absolute transcript abundance, dictates downstream developmental outputs. Constitutive overexpression of *35S::CLV3* constitutively represses *WUS* and terminates stem cell activity, arresting both organogenesis and stem elongation as previously reported ^50^. By contrast, *pTOC1::CLV3*, which recapitulates endogenous circadian *CLV3* oscillation, fully rescues all developmental defects of *clv3-1*, restoring normal SAM morphology, organ production and diurnal stem growth rhythms (Fig. 3a-3c). Under diurnal cycles, dusk-enriched TOC1 elevates CLV3 levels to moderately constrain WUS activity, while reduced CLV3 signaling at dawn permits moderate WUS accumulation and nocturnal stem elongation (Fig. 5 and Extended Data Fig. 5), partitioning organ initiation and stem growth into distinct time windows. Considering the 24-h circadian expression of *CLV3* and 12-h rhythmic expression of *WUS* but near 24-h accumulation of WUS in which the maximum *CLV3* expression is at ZT12 but represses *WUS* at ZT0-ZT12 (Fig. 5 and 6), there should be dynamic and intricate regulations between *CLV3*-*WUS* module at temporally transcriptional, translational and post-translational levels. Moreover, cell-type-specific responses to identical *CLV3*-*WUS* oscillatory signals remain elusive. The same circadian fluctuation of WUS and CLV3 simultaneously controls stem cell fate in the CZ and cell elongation in the RM, yet these two meristematic domains exhibit divergent transcriptional programs ^63^. Cell-specific co-factors, histone modifications or DNA methylation landscapes that shape differential target gene activation downstream of WUS require further exploration.

WUS acts as a hub gene of complicated gene regulatory networks in meristem maintenance and cell proliferation^51,59^. Recent studies revealed that PMEs-mediated pectin metabolism in cell wall organization that was controlled by WUS in SAM was critical for stem cell identity, meristem maintenance and organ initiation and growth ^57^. Region-specific *PME* expression creates mechanical partitioning within the SAM to separate stem cell maintenance and organ/stem outgrowth ^43^. We found here that WUS directly binds the G-box element within the *PME5* promoter to activate its transcription. Overexpression of *PME5* partially rescues the shortened RM and dwarf stem phenotypes of *wus-7* mutants, demonstrating PME activity is sufficient to restore stem growth downstream of WUS. Therefore, diurnal WUS concentration gradients originating from the OC lead to oscillatory PME activation in the distant RM to drive stem elongation (Fig. 7), while high WUS abundance within the CZ maintains hyper-methylated rigid cell walls that restrict stem cell overexpansion ^57^, spatially segregating meristem maintenance and growth functions. Moreover, WUS served as the genetic foundation mediating the effects of AP1, TOC1 and CLV3 on stem development (Fig. 6). Therefore, combined with previous findings ^36,57^, we dissected a spatiotemporal regulatory cascade, in which the *AP1*/*CCA1*-*TOC1* module modulated *CLV3*-WUS-*PMEs* dynamics to gate stem development and organogenesis (Extended Data Fig. 7): The FM-localized AP1/CCA1 complex acts as a tissue-specific timer to sustain rhythmic *TOC1* expression. Mobile TOC proteins transmit temporal signals non-cell-autonomously from floral primordia to IMs, directly driving oscillatory *CLV3* transcription and conferring circadian fluctuations to WUS protein abundance and subsequent *PMEs* expression. This creates a tissue-crossing timing signal that links floral identity programs with meristem growth dynamics to pattern optimal plant shoot morphology.

## METHODS

### Plant materials and chemical treatments

#### Arabidopsis thaliana

ecotype Landsberg *erecta* (L*er*) was used as the wild type (WT) reference in this study. The following previously characterized Arabidopsis lines were also used: *clv3-1* ^64^, *wus-7* ^65^, *pWUS::WUS-GFP;wus-101* ^66^. All *Arabidopsis thaliana* plants were cultured either in a greenhouse or in a growth chamber (CU-36L4, Percival Scientific; BPC600H, JIUPO, China) maintained at 22°C under a 12 h light/12 h dark photoperiod (ZT0-ZT12 (light: 09:00-21:00); ZT12-ZT24 (dark: 21:00-09:00)) or constant light (LL) condition. Five-week-old *Nicotiana benthamiana* plants cultivated in soil under identical growth conditions were used for Agrobacterium-mediated transient transformation.

For peptide and chemical treatments, CLV3 peptide was dissolved in ddH_2_O to a final stock concentration of 10 µM. Dexamethasone (DEX) was first dissolved in DMSO to prepare a 10 mM stock solution and further diluted with ddH_2_O to 10 µM. Before application, 0.1% (v/v) Silwet L-77 was added to each working solution. Solutions were directly applied onto SAMs.

#### Plasmid construction

The promoter and coding sequences (CDS) of *TOC1*, *CCA1*, *AP1*, *WUS* and *CLV3* were amplified using WT genomic DNA or cDNA as the template. The resulting fragments were then cloned into the pMDC107, pMDC83 and pEarleyGate301 vectors via restriction digestion and ligation to generate the following in-frame fusion constructs: *pTOC1::CLV3*, *pCCA1::CLV3*, *pWUS::WUS*, *pAP1::TOC1*, and *pCLV3::CLV3-3’UTR*, respectively. To generate mutated versions of *pTOC1::CLV3*, site directed mutagenesis and PCR amplification was performed to obtain *pTOC1_EEm_::CLV3*, *pTOC1_CArGm_::CLV3* and *pTOC1_CArGmEEm_::CLV3.* And to generate a mutated version of *pCLV3::CLV3-3’UTR*, we also site directed mutagenesis and PCR amplification was performed to obtain *pCLV3_EEm1_::CLV3-3’UTR_EEm2_.* In addition, the CDS of *TOC1*, *WUS*, and *CLV3* were amplified using WT cDNA as a template and cloned into pMDC83 vector to generate the *p35S::CLV3*, *p35S::TOC1* and *p35S::WUS* construct. To generate the *p35S::TOC1-GR-3FLAG* construct, the *TOC1* coding region was amplified and fused at the C-terminus with a GR (glucocorticoid receptor) domain and three copies of FLAG epitope tag (3FLAG, DYKDDDDKDYKDDDDKDYKDDDDK), and the resulting version was inserted into the pEarleyGate100 vector. To construct *pCLV3::LUC-3’UTR*, a 5 kb sequence upstream of the *CLV3* start codon and a 2 kb sequence downstream of the stop codon were amplified and cloned into the pEarleyGate301 vector, followed by the insertion of the LUC reporter gene downstream of the *CLV3* coding region. Site directed mutagenesis of this construct (*pCLV3:LUC-3’UTR*) was then performed to generate *pCLV3_EEm1_::LUC-3’UTR*, *pCLV3::LUC-3’UTR_EEm2_* and *pCLV3_EEm1_::LUC-3’UTR_EEm2_*.

In addition, the promoter of *PME5* was amplified using WT DNA as a template and cloned into pEarleyGate301 vector, followed by the insertion of the LUC reporter gene downstream of the *PME5* coding region to construct *pPME5::LUC*, and then site directed mutagenesis of this construct to generate *pPME5_G-boxm_::LUC*. All constructs were introduced into *Agrobacterium tumefaciens strain GV3101* for downstream assays. All constructs were sequence-verified. Primers are provided in Extended Data Table 1.3.

#### Histological sectioning

For semithin section, the inflorescences of *Arabidopsis* were fixed using FAA solution (50% ethanol, 5% glacial acetic acid, and 3.7% formaldehyde). Samples were then subjected to vacuum infiltration for 30 minutes to enhance fixation, followed by graded ethanol dehydration (30%, 50%, 70%, 95%, and 100%, 30 minutes each step). Safranin O (0.05%) was added to the 95% and 100% ethanol solutions to stain the tissues. Following dehydration, samples were gradually infiltrated and embedded in resin blocks using a resin embedding kit (Kulzer Technik, Technovit 7100) according to the manufactureŕs instructions. Embedded tissues were sectioned to a thickness of 1µm for microscopic analysis.

For paraffin sectioning, aboveground tissues from 13-day-old *Arabidopsis* seedlings were harvested and fixed in FAA solution (50% ethanol, 5% glacial acetic acid, and 3.7% formaldehyde) under vacuum infiltration for 30 minutes, followed by fixation at room temperature for an additional 24 hours. Samples were then dehydrated through a graded ethanol series (50%, 70%, 95%, and 100% ethanol, 30 minutes per step). Tissue infiltration was performed using a xylene substitute (Histoclear) in increasing concentrations: 30% Histoclear/70% ethanol, 65% Histoclear/35% ethanol, and 100% Histoclear (30 minutes each step). Following complete infiltration, tissues were embedded in paraffin. Embedded samples were sectioned at a thickness of 6 µm using a rotary microtome (Leica RM2235, Leica, Shangai).

#### Confocal microscopy

The inflorescences of Arabidopsis were excised, and the mature floral organs along with surrounding primordia adjacent to the SAM were carefully removed using a fine needle under a stereomicroscope. The exposed meristems were stained with 50 μg/mL propidium iodide (PI, Sigma-Aldrich) for 10 minutes to visualize cell walls, then mounted on grooved slides for imaging. Fluorescence imaging was performed using a Leica Stellaris 5 confocal laser scanning microscope. Imaging was performed sequentially through GFP (excitation 488 nm/emission 507 nm) and PI (excitation 493 nm/emission 636 nm) channels, supplemented with additional bright-field observations when required. Red channel signals and bright-field images were converted to grayscale conversion in LAS X software prior to fusion with green channel signals (GFP). Identical fluorescence intensity settings were maintained across experimental replicates. Imaging parameters included the use of a 20× objective lens (numerical aperture (NA) 0.75), and a Z-step size of 2 µm.

#### Scanning Electron Microscopy

Fresh inflorescence SAM tissues of Arabidopsis were rapidly dissected under a stereomicroscope without chemical fixation or dehydration. Dissected samples were mounted onto sample stubs using conductive carbon double-sided tape and loaded onto the Peltier cold stage (Deben MK3 Coolstage) of a Hitachi TM3030Plus tabletop scanning electron microscope. After chamber evacuation, the cold-stage temperature was set to-20 °C. Secondary-electron images were collected. At least 10 independent biological samples were examined for each genotype, and representative micrographs are presented.

#### Confocal microscopy

To observe fluorescence signals in SAMs, bolted inflorescence apices were dissected under a stereomicroscope; mature floral organs and surrounding primordia were carefully removed with fine needles. Dissected samples were stained with 50 µg mL^-1^ propidium iodide (PI; Sigma-Aldrich) for 10 min and mounted on grooved glass slides. Confocal imaging was performed on a Leica Stellaris 5 microscope. GFP signals were collected at excitation 488 nm/emission 507 nm; PI signals were collected at excitation 493 nm/emission 636 nm. Bright-field images were acquired when necessary. Red-channel and bright-field images were converted to grayscale in LAS-X software before merging with green-channel GFP signals. Identical imaging parameters were kept across all biological replicates. Images were obtained using a 20 × objective lens (NA = 0.75), with a Z-step size of 2 µm.

#### RNA extraction and quantitative RT-PCR

SAMs were manually dissected from plants. Sample collection and treatment, as well as the timing, were performed as previously described by our laboratory ^36^. Total RNA was extracted using RNAiso Plus (TaKaRa). Genomic DNA contamination was eliminated by DNase I (Roche) digestion. First-strand cDNA was synthesized with the RevertAid First-Strand cDNA Synthesis Kit (TaKaRa). Quantitative RT-PCR (RT-qPCR) was performed with SYBR Green qPCR Master Mix (Vazyme) on a Bio-Rad CFX-96 real-time PCR system. Three independent biological replicates were assayed. Gene expression levels were normalized against *IPP2* as the internal reference. All RT-qPCR primer sequences are provided in Extended Data Table 1.3.

#### Chromatin immunoprecipitation (ChIP) assay

ChIP experiments were performed as previously described ^67^. SAM tissues (∼2g) were harvested from bolted L*er* and *pTOC1::TOC1-HTF* transgenic plants. For WUS ChIP assays, SAM tissues were collected from Ler and *pWUS::WUS-GFP* transgenic plants. Tissues were ground to fine powder in liquid nitrogen and homogenized in M1 buffer (10 mM phosphate buffer pH 7.0, 0.1 M NaCl, 10 mM β-mercaptoethanol, 1 M hexylene glycol, protease-inhibitor cocktail, 1 mM PMSF). Cross-linking was done with 37% formaldehyde for 10 min, and homogenates were filtered through Miracloth. Nuclei were pelleted by centrifugation at 2000 × g. Pellets were sequentially washed with M2 buffer (M1 buffer supplemented with 10 mM MgCl_2_ and 0.5% Triton X-100) and M3 buffer (M1 buffer without 1 M hexylene glycol). Nuclei were resuspended in nuclear-lysis buffer (50 mM Tris-HCl pH 8.0, 10 mM EDTA, 1% SDS, protease-inhibitor cocktail). Chromatin was sonicated to generate DNA fragments of 200-500 bp. Immunoprecipitation was carried out using anti-FLAG antibody or anti-GFP antibody. Purified immunoprecipitated DNA was used for ChIP-qPCR or ChIP-seq library construction. ChIP-qPCR primer sequences are listed in Extended Data Table 1.3.

#### Stem-growth rate measurement

Plants were acclimated for one week under 12 h light /12 h dark cycles (light: 09:00-21:00; dark: 21:00-09:00). A thin bamboo stake was placed adjacent to each individual plant. Stem height was marked on the stake at 09:00 and 21:00 every day. Day-and night-time stem elongation values were calculated by measuring the distance between successive marks. Silique number per unit length was determined by counting siliques in the 2 cm segment above the first silique on the main inflorescence stem of Arabidops.

#### Luciferase complementation imaging (LCI) assay

Reporter constructs including *pCLV3::LUC-3’UTR*, *pCLV3_EEm1_::LUC-3’UTR*, *pCLV3::LUC-3’UTR_EEm2_*, *pCLV3_EEm1_::LUC-3’UTR_EEm2_*, pPME5::LUC, *pPME5_G-boxm_::LUC*, and effector constructs *p35S::TOC1-GFP*, *p35S::TOC1-GR-3FLAG*, *p35S::WUS-GFP* were transformed into *Agrobacterium tumefaciens* strain GV3101. Agrobacterial cultures carrying reporter and effector plasmids were mixed at appropriate ratios and infiltrated into the abaxial side of *N. benthamiana* leaves. Infiltrated plants were kept in darkness for 10 h and then grown under light conditions for 48 h. Firefly luciferase substrate buffer was sprayed onto leaf surfaces before luminescence imaging, as described previously ^36^.

#### *In situ* hybridization

*In situ* hybridization was performed as previously described ^67^. Arabidopsis inflorescences were collected and fixed in 4% (w/v) paraformaldehyde (PFA), dehydrated through a graded ethanol series, and embedded in paraffin. Tissue sections were cut at a thickness of 8 µm for in situ hybridization. RNA probe templates were amplified from cDNA using gene-specific primers containing T7 or T3 promoter sequences at their 5′ ends. Digoxigenin (DIG)-labeled RNA probes were synthesized *in vitro* using T7/T3 RNA polymerase (Roche, 10881767001) and digoxigenin-UTP (Roche, 11277073910).

#### Protein extraction and immunoblot analysis

Dissected SAM tissues were ground to powder in liquid nitrogen inside 1.5 mL tubes. Tissue powder was mixed with 100 µL of 2 × SDS loading buffer (1 M Tris-HCl pH 6.8, 10% glycerol, 4% SDS, 0.005% bromophenol blue, 10% β-mercaptoethanol). Samples were heated at 95 °C for 10 min and centrifuged at 12 000 × g for 5 min. Supernatants were separated by 10% SDS-PAGE and electro-transferred onto polyvinylidene difluoride (PVDF) membranes (Immobilon-P; Millipore). Membranes were blocked with 5% skim-milk solution for 1 h, then incubated with primary antibodies (anti-GFP, anti-Actin) for 4 h. After three washing steps, membranes were incubated with HRP-conjugated anti-mouse secondary antibody (1:5000; Agrisera, AS09602), followed by three additional washes. Target proteins on PVDF membranes were detected by enhanced chemiluminescence (ECL) using Super Signal West Femto Maximum-Sensitivity Substrate (Thermo Scientific) on a ChemiDoc Touch imaging system (Bio-Rad). Time-course WUS-GFP protein abundance assays were repeated for at least three independent biological replicates.

#### Accession Numbers

Sequence data for the genes in this article can be found in the Arabidopsis Genome Initiative or GenBank/EMBL databases under the following accession numbers: AP1: AT1G69120, CAL: AT1G26310, CCA1: AT2G46830, CLV3: AT2G27250, IPP2: AT3G02780, PME5: AT5G47500, TOC1: AT5G61380, VGD1: AT2G47040, WUS: AT2G17950.

## SUPPLEMENTAL INFORMATION TITLES AND LEGENDS

**Extended Data Fig.1.**
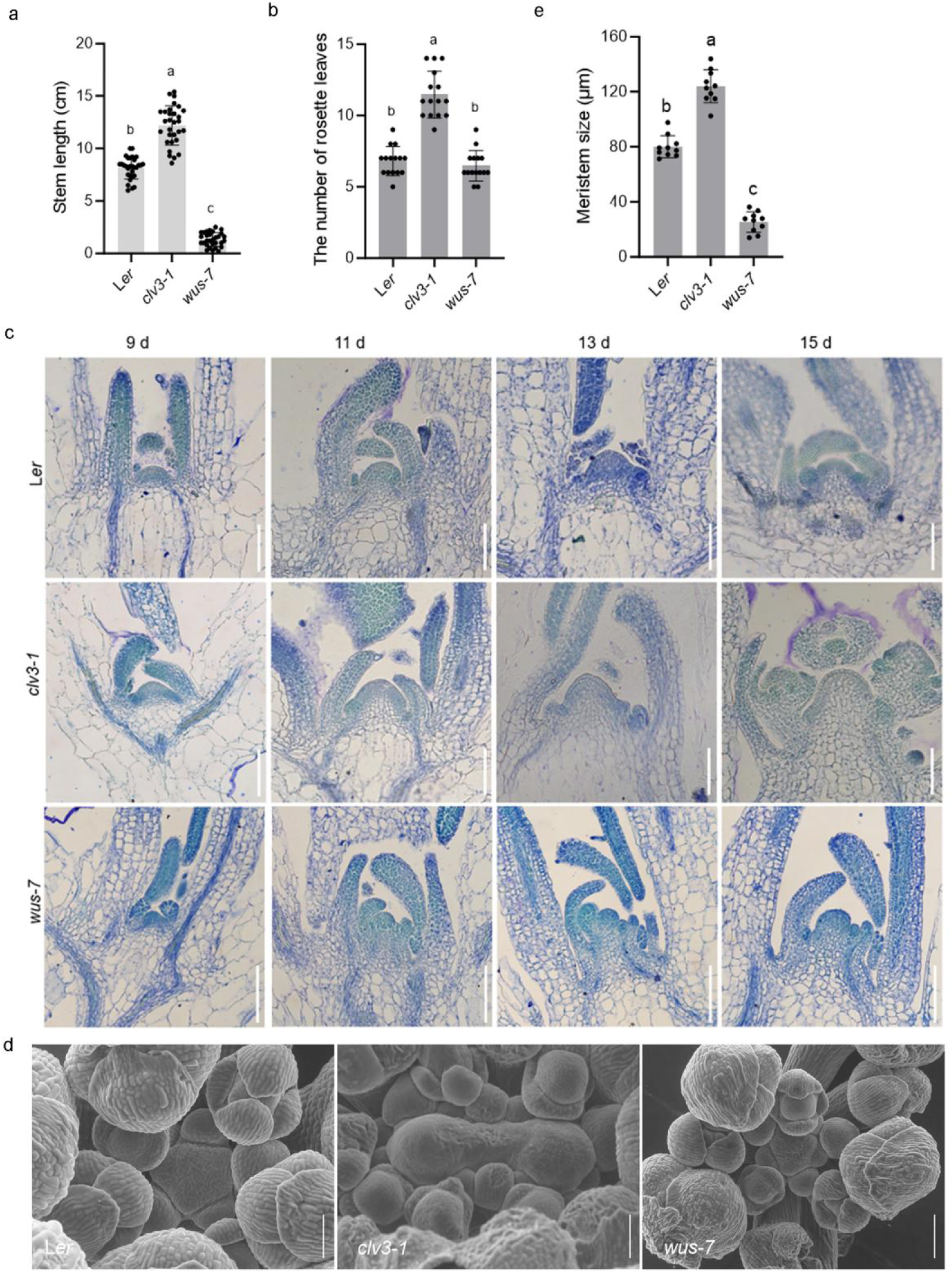
CLV3 and WUS control stem development and SAM activity. **(a)** Statistical analysis of the length of stem of indicated plants. Data represent mean ± SD (*n* = 30). Different letters indicate statistically significant differences by one-way ANOVA (Tukey’s multiple comparisons test, *P* < 0.01). **(b)** Number of rosette leaves of indicated plants. Data represent mean ± SD (*n* = 15). Different letters indicate statistically significant differences by one-way ANOVA (Tukey’s multiple comparisons test, *P* < 0.01). **(c)** Histological sectioning of shoot apices at 9, 11, 13, and 15 days after germination (DAG) of indicated plants. Scale bars = 50 μm. **(d)** Representative scanning electron microscopy (SEM) images of SAMs from L*er*, *clv3-1* and *wus-7*. Scale bars = 50 μm. **(e)** Quantification of SAM size of indicated plants. Data are mean ± SD (*n* = 10). Different letters indicate statistically significant differences by one-way ANOVA (Tukey’s multiple comparisons test, *P* < 0.01).

**Extended Data Fig.2.**
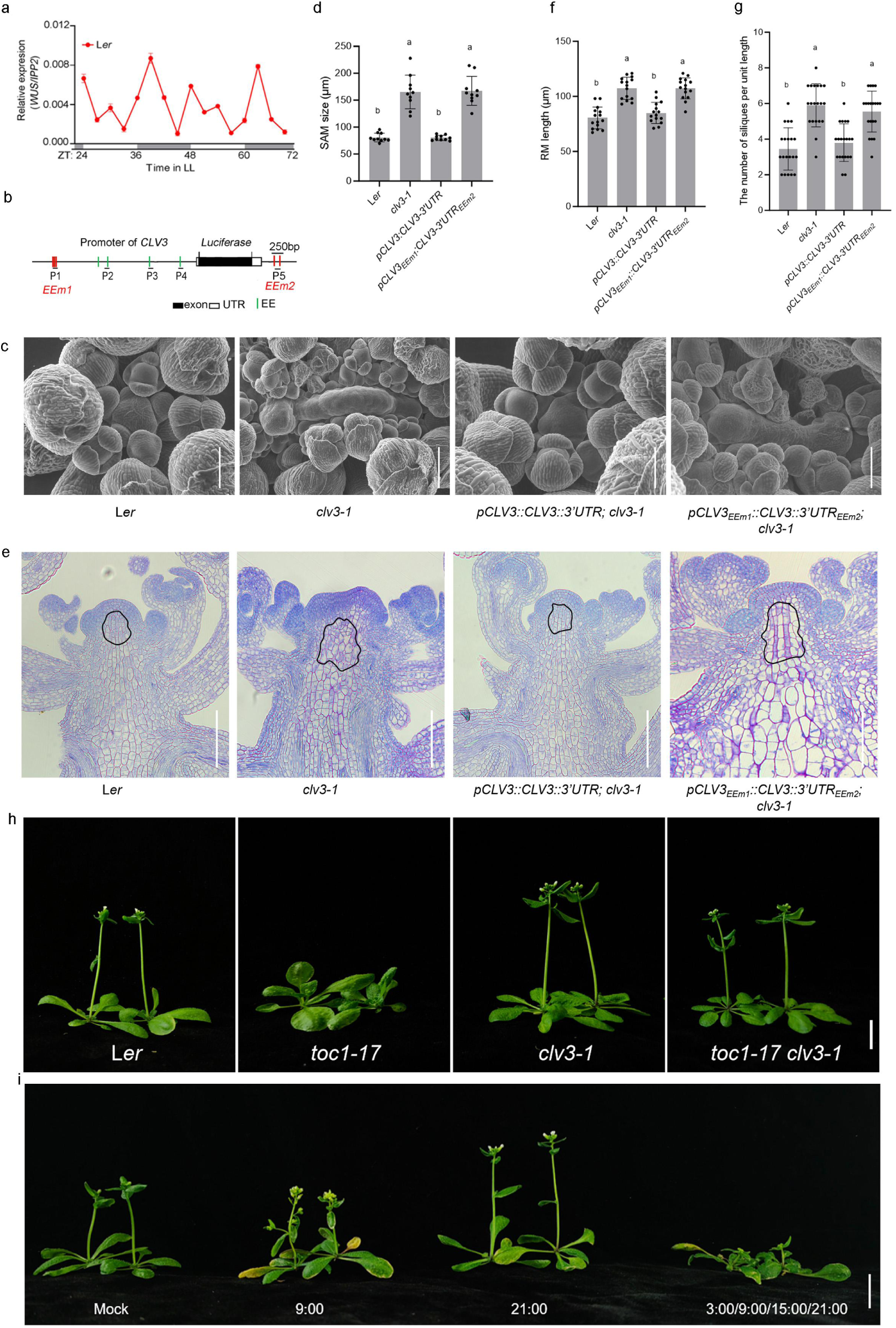
TOC1 activates *CLV3* to maintain meristem activity and stem development. **(a)** RT-qPCR to examine the expression level of *WUS* in L*er*. Data represent mean ± SD of three biological replicates. *IPP2* was used as a normalization control. White and gray bars represent subjective day and night, respectively. **(b)** Diagram of the *pCLV3::LUC-3’UTR* construct. The coding region of *CLV3* was replaced with the *LUC* gene and the mutated EE motifs were indicated with red lines designated EEm1 and EEm2, respectively. **(c)** Representative scanning electron microscopy (SEM) images of SAMs of indicated plants. Scale bars = 100 μm. **(d)** Size of the SAM of indicated plants. Data are mean ± SD (*n* = 10). Different letters indicate statistically significant differences by one-way ANOVA (Tukey’s multiple comparisons test, *P* < 0.01). **(e)** Longitudinal sections of shoot apices of indicated plants showing the length of RM. Scale bars = 100 μm. **(f)** Statistical analysis of the length of RM of indicated plants. Data represent mean ± SD (*n* = 15). Different letters indicate statistically significant differences by one-way ANOVA (Tukey’s multiple comparisons test, *P* < 0.01). **(g)** Statistical analysis of the length of siliques per unit length of indicated plants. One unit length=2 cm. Data represent mean ± SD (*n* = 15). Different letters indicate statistically significant differences by one-way ANOVA (Tukey’s multiple comparisons test, *P* < 0.01). **(h)** Whole-plant phenotypes of indicated plants. Scale bars: 1 cm. **(i)** Whole-plant phenotypes of *toc1-17* plants treated with CLV3-peptide for 7 days at the indicated time points. Plants were grown under 12/12 (linght/dark) conditions for an additional 10 days after treatment before imaging. Scale bars: 1.5 cm.

**Extended Data Fig.3.**
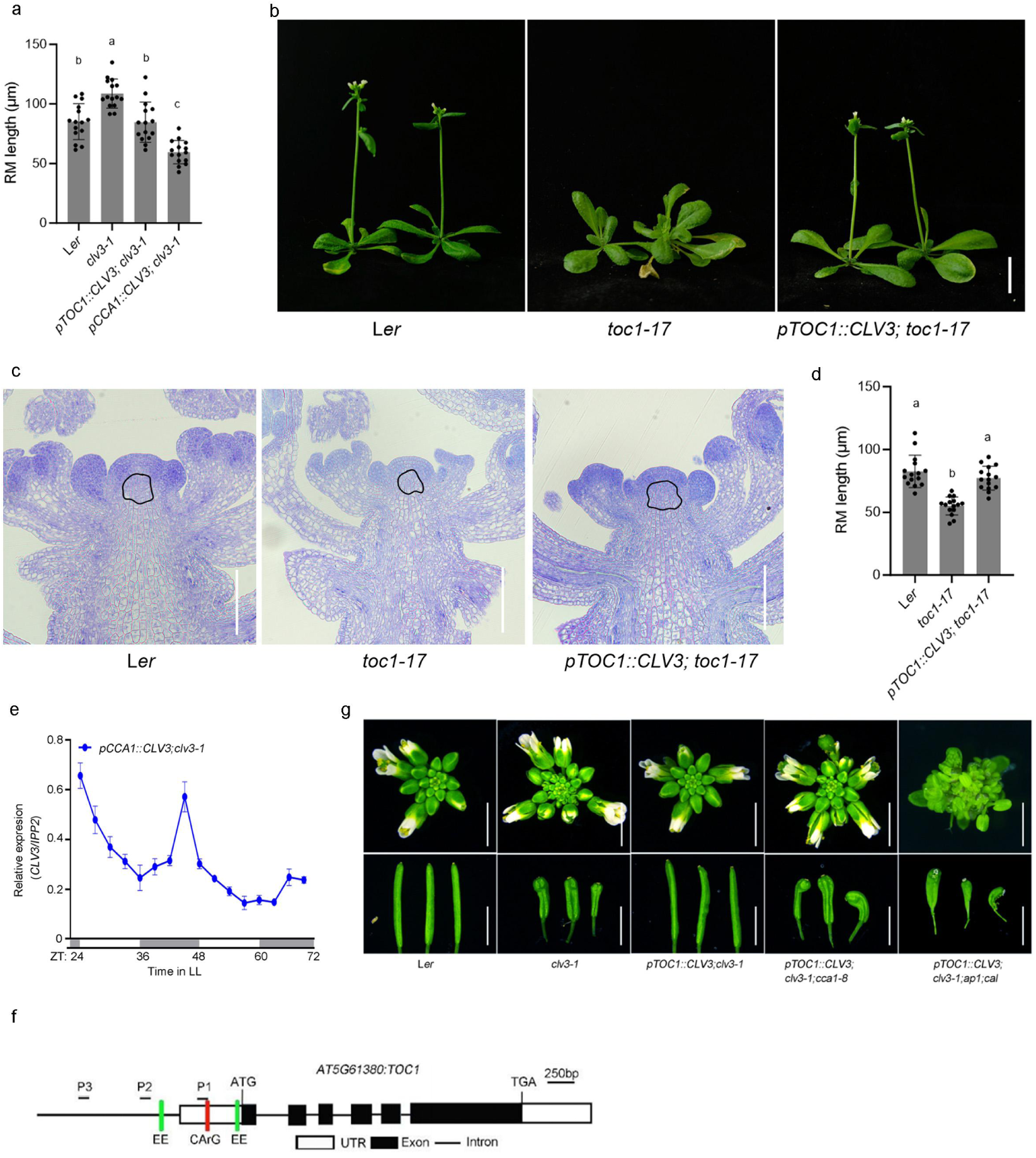
The circadian expression pattern of CLV3 determines SAM activity and stem elongation. **(a)** Statistical analysis of the length of RM of indicated plants. Data represent mean ± SD (*n* = 15). Different letters indicate statistically significant differences by one-way ANOVA (Tukey’s multiple comparisons test, *P* < 0.01). **(b)** The whole-plant phenotypes of indicated plants. Scale bars: 1 cm. **(c)** Longitudinal sections of shoot apices of indicated plants showing the length of RM. Scale bars = 150 μm. **(d)** Statistical analysis of the length of RM of indicated plants. Data represent mean ± SD (*n* = 15). Different letters indicate statistically significant differences by one-way ANOVA (Tukey’s multiple comparisons test, *P* < 0.01). **(e)** RT-qPCR to examine the expression of *CLV3* in *pCCA1::CLV3;clv3-1*. Data represent mean ± SD of three biological replicates. *IPP2* was used as a normalization control. White and gray bars represent subjective day and night, respectively. **(f)** Diagram of the *TOC1* locus. White and black rectangles and black lines represent untranslated regions, coding regions, and introns, respectively. Green lines and red line represent EE motifs and CArG. **(g)** Phenotypes of inflorescences and siliques of indicated plants. Scale bars: 0.5 cm.

**Extended Data Fig.4.**
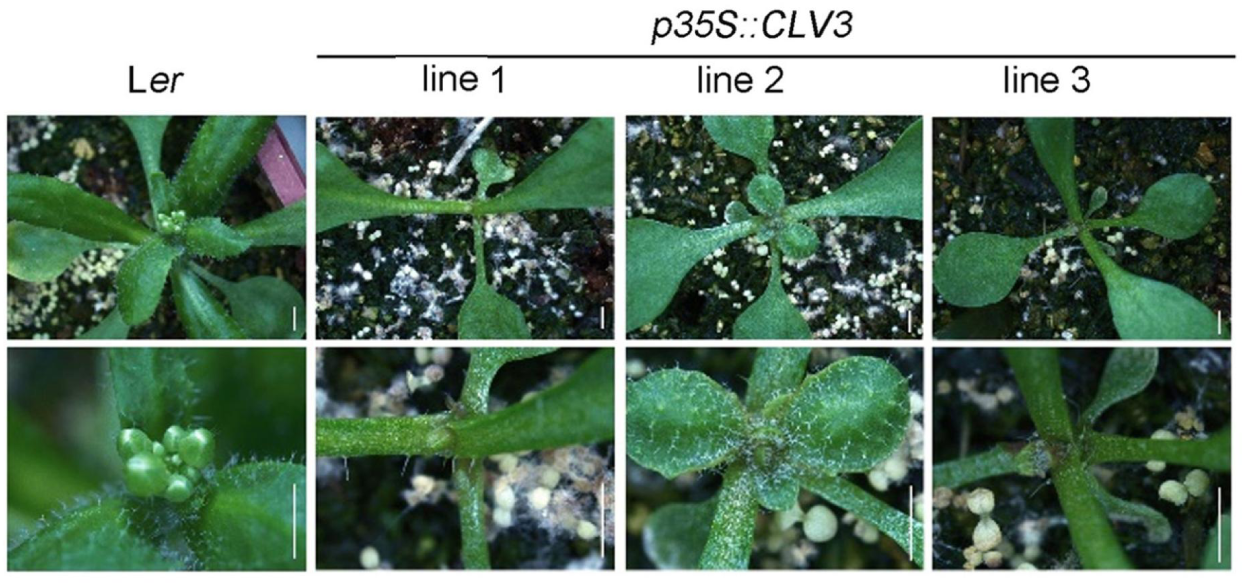
Phenotypes of *CLV3* overexpression lines. Morphological observation of the L*er* and different *p35S::CLV3* transgenic lines. Scale bars: 0.5 cm.

**Extended Data Fig.5.**
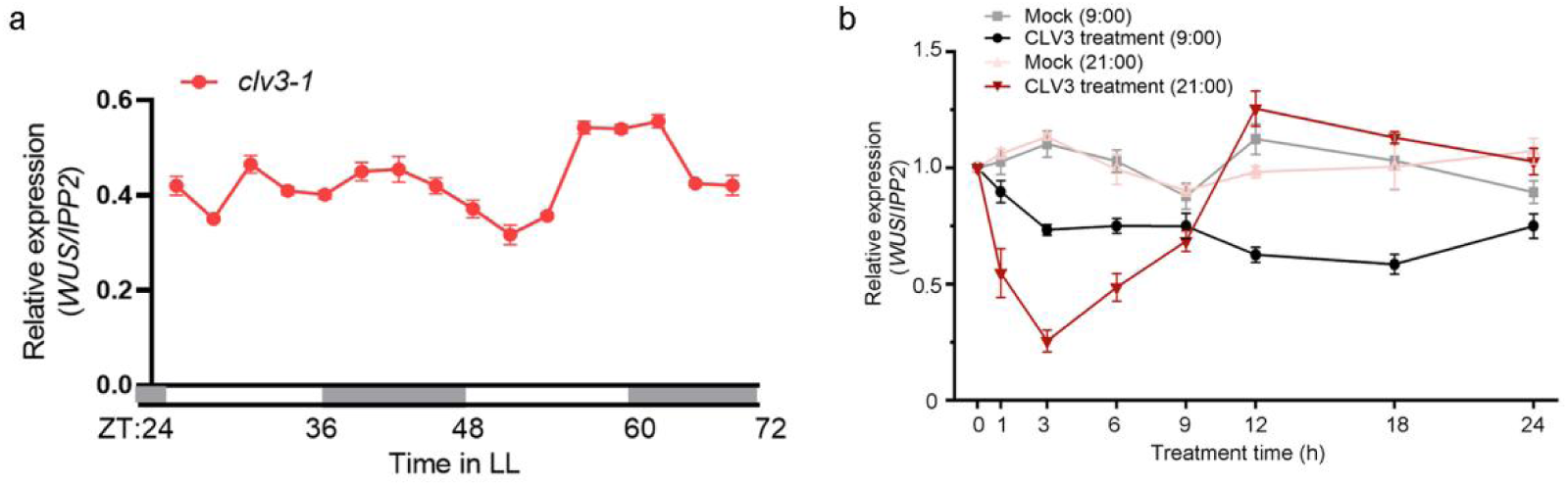
CLV3 is required for the rhythmic expression of *WUS*. **(a)** RT-qPCR to examine the expression of *WUS* in *clv3-1*. Data represent mean ± SD of three biological replicates. *IPP2* was used as a normalization control. White and gray bars represent subjective day and night, respectively. **(b)** RT-qPCR to examine the expression of *WUS* at the indicated time points after single CLV3-peptide treatment of *clv3-1* plants at ZT0 (9:00) and ZT12 (21:00), respectively. *IPP2* was used as a normalization control. *WUS* expression at timepoint 0 was set as 1. Data represent mean ± SD of three biological replicates.

**Extended Data Fig.6.**
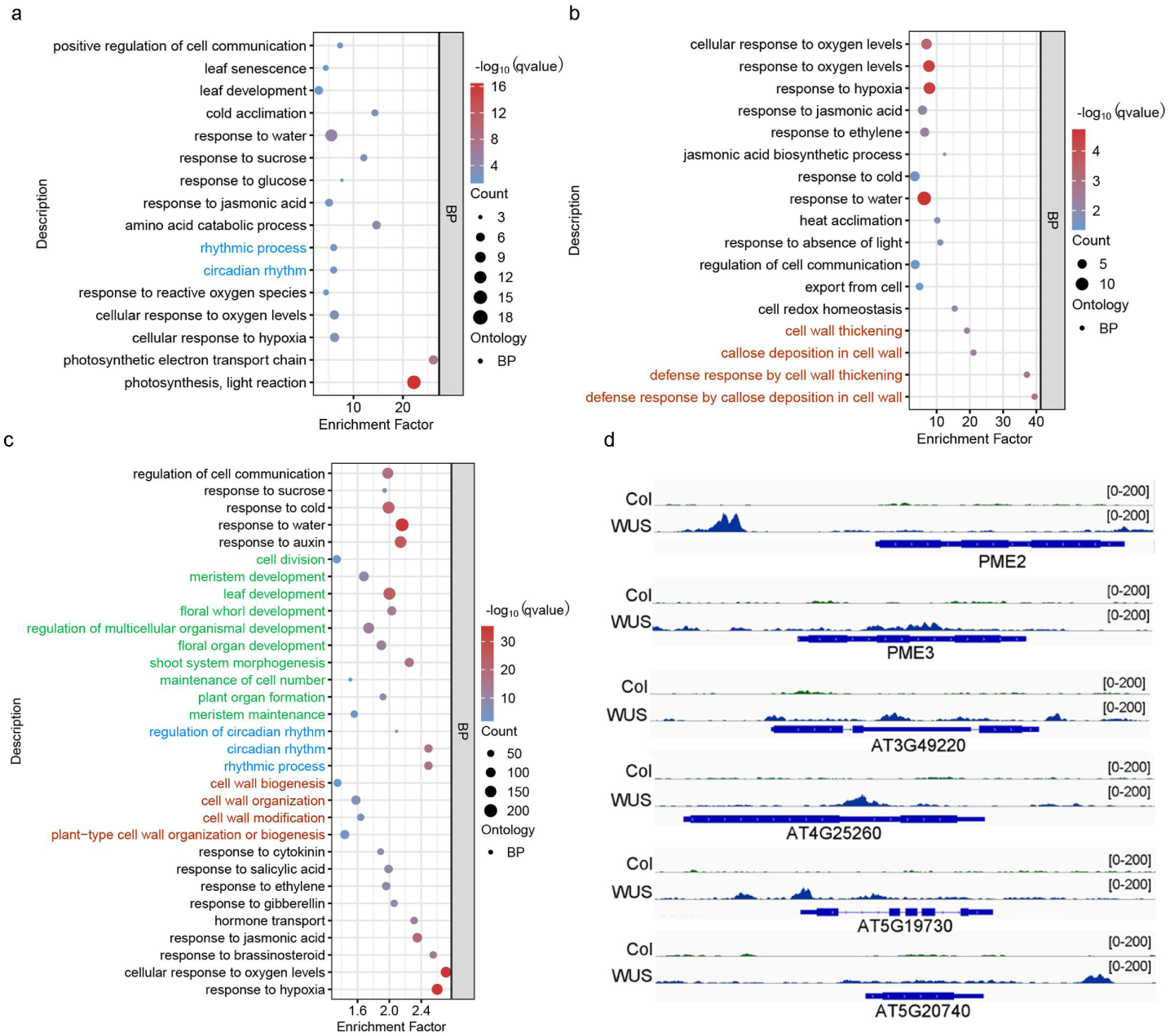
Omics analysis of WUS-regulated genes. **(a)** RNA-seq analysis of differentially expressed genes from *35S::WUS-GR* plants. Gene Ontology (GO) enrichment showing that WUS-regulated genes. (b) GO enrichment analysis of WUS-repressed genes. (c) GO analysis of WUS-binding loci identified by ChIP-seq. (d) Identification of *PMEs* directly bound by WUS. Several *PME* genes were isolated as direct WUS targets.

**Extended Data Fig.7.**
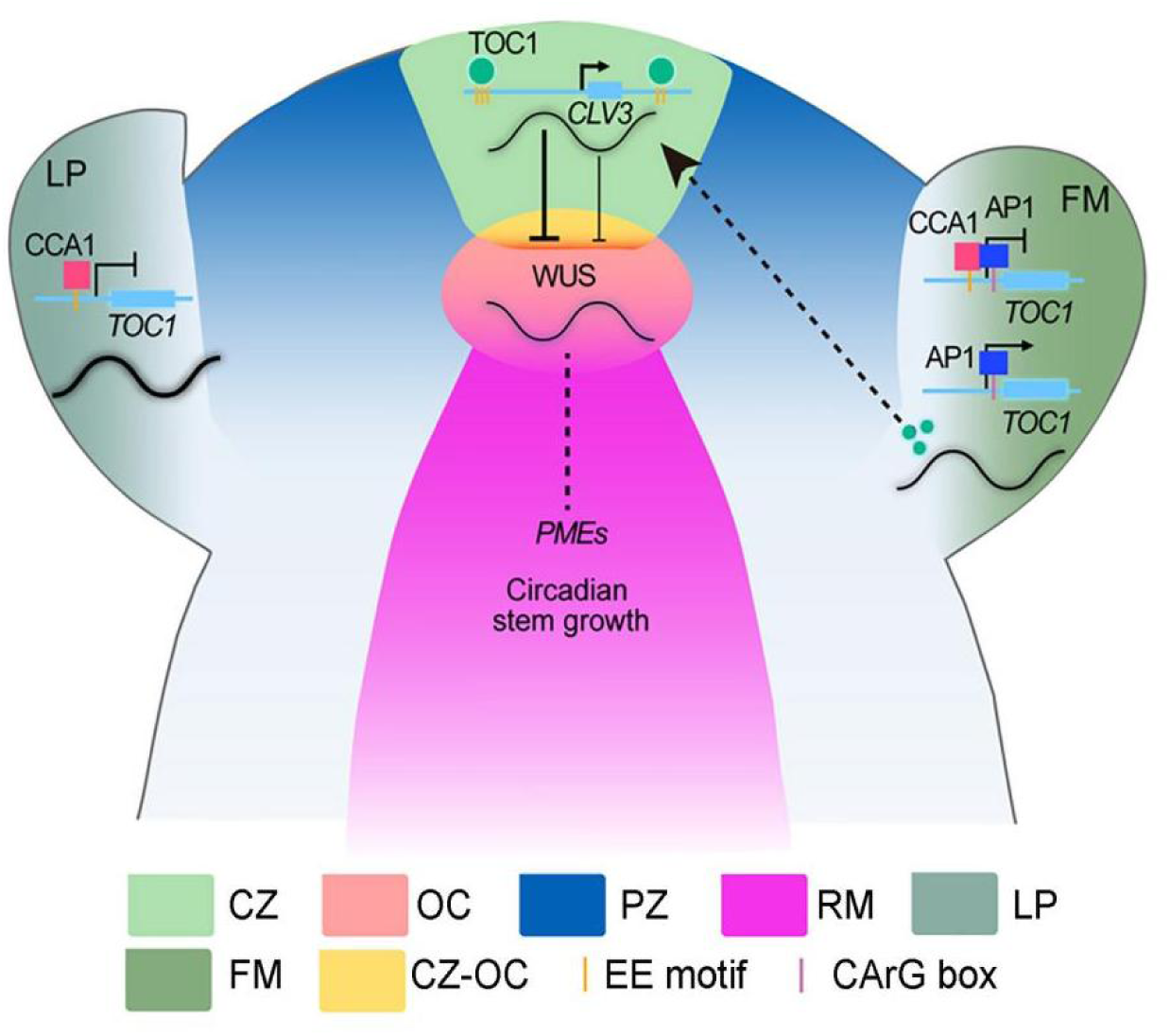
The work model: A spatiotemporal cascade of regulatory events that gates stem development. AP1 and CCA1 maintain both the expression level and rhythmicity of *TOC1* within the FM primordium. This regulation non-cell-autonomously and spatially activates RMs and initiates stem development. Subsequently, TOC1 directly activates *CLV3* expression to maintain its circadian expression, which is critical for the circadian accumulation of WUS, and ultimately, for temporally circadian-regulated stem growth.

**Extended Data Table 1.1.** Differentially Expressed Genes (DEGs) were identified between *WUS-GR* (DEX) compared to Col (DEX).

**Extended Data Table 1.2.** WUS binds genes.

**Extended Data Table 1.3.** Primers list in this work.

## ACKNOWLEDGMENTS

We thank Dr. Robert Sablowski, Wenhao Yan, Xuemei Chen, Yuling Jiao, and Zhong Zhao for the valuable discussion and advice. This work was supported by the National Natural Science Foundation of China (grants U24A20391, 32270340 and 32670445 to X.L., 32671106 to H.Z.), Hebei Natural Science Foundation (grants C2025205068 to X.L., and C2024205009 to H.Z.), and the Seed Science and Technology Innovation Team Project of Shijiazhuang (232490472A to X.L.), Hebei Education Department (grant JCZX2025026 to X.L.), startup funds from Hebei Normal University (L2021B19 to X.L., and L2024B19 to H.Z.).

## AUTHOR CONTRIBUTIONS

X.L. and H.Z. conceived the project, designed the experiments, and wrote the manuscript. J.Q., J.X. and H.Z. performed tissue sampling, gene expression analysis such as RT-qPCR and *in situ* hybridization, Confocal analysis and histological section.

Y.Y. performed the RNA-seq analysis and ChIP-seq analysis under the directions of X.X. and Q.X.. J.Q., M.J., and Y.Z. performed genetic and transgenic experiments, phenotypic analyses, protein-protein interaction, J.Q. analyzed and discussed the results and wrote the manuscript with X.L. and H.Z.

